# *XIST* Drives X-Chromosome Inactivation and Safeguards Female Extraembryonic Cells in Humans

**DOI:** 10.1101/2025.11.19.689206

**Authors:** Amitesh Panda, Léo Carrillo, Bradley Philip Balaton, Jeanne Brouillet, Solomon Nshemereirwe, Jarne Bonroy, Charbel Alfeghaly, Romina Facchinello, Sherif Khodeer, Nicolas Peredo, Ruben Boers, Gael Castel, Charlie London, Emmanuel Cazottes, Madeleine Moscatelli, Raissa Songwa Tchinda, Thi Xuan Ai Pham, San Kit To, Ryan Nicolaas Allsop, Yang Wang, Desislava Staneva, Peter J. Rugg-Gunn, Kathy K. Niakan, Joost Gribnau, Jean-François Ouimette, Claire Rougeulle, Vincent Pasque

## Abstract

Dosage compensation of sex chromosomes through X-chromosome inactivation (XCI) is required for mice extra-embryonic tissue growth and embryo development. The species specificity in mechanisms and timing leading to XCI during early embryogenesis, however, left the key question of the interdependence between XCI and human development open. Here, we show that the differentiation of naive human pluripotent stem cells to trophoblast stem cells and extraembryonic mesoderm cells triggers XCI. The inactive X chromosome, however, displays an atypical chromatin state, lacking classical enrichment of heterochromatin markers and DNA methylation. We demonstrate that extraembryonic differentiation and XCI are kinetically and functionally linked. Using loss of function approaches, we prove that *XIST* is required for human XCI establishment. We also reveal that XCI is key for the survival of human female extraembryonic cells. Our work therefore links XCI to the formation of extraembryonic annexes, with important consequences for human reproductive biology.

**HIGHLIGHTS:**

- Naive hPSCs to EXMCs and TSCs differentiation recapitulates human XCI
- The Xi has an unusual chromatin status in human extraembryonic cells
- *XIST* is required for the establishment of human XCI
- XCI supports healthy development of human female extraembryonic cells

## INTRODUCTION

X-chromosome inactivation (XCI) is the process that equalizes X-linked gene dosage between XX female and XY male eutherian mammals through the silencing of one of the two X chromosomes in females ^1^. In mice, XCI is triggered by the long non-coding RNA *Xist*, which recruits co-factors for transcriptional silencing, including modifiers of RNA and chromatin, leading to the formation of a heterochromatic inactive X chromosome (Xi) ^2–8^. XCI is established during early mice embryonic development, first in an imprinted manner where it is always the paternal X chromosome that is inactivated in female preimplantation embryos and in extraembryonic tissues ^5,6,9–12^. Impairing XCI establishment through *Xist* mutation results in early female lethality, which has been attributed to defects in extraembryonic tissues in general, and in extraembryonic ectoderm placental progenitors in particular. The importance of *Xist* and XCI for extraembryonic development has therefore been abundantly documented in mice ^13–15^.

In contrast, the kinetics, mechanisms, and importance of XCI for the development of human extraembryonic annexes remain poorly understood, and the role of *XIST* in these processes has never been addressed, despite potential consequences for embryo implantation, pregnancy success, and sex-biased diseases of developmental origin ^16^. This is all the more relevant as major differences exist between rodents and humans, both in terms of X chromosome status and developmental features ^17^. Indeed, in humans, *XIST* is expressed from the two X chromosomes during female preimplantation development without triggering XCI ^18,19^. Petropoulos and colleagues suggested that instead of classical XCI, X-linked gene expression is transiently and partially reduced or ‘dampened’ from both active X chromosomes in the preimplantation human embryo, a stage during which *XIST* accumulates on the X chromosomes in a dispersed configuration ^19–21^. Despite initial debates over the observation of dampening ^19,20,22^ various reports have now provided evidence of X-chromosome dampening in human primordial germ cells, pre-implantation human embryos and in naive human pluripotent stem cells ^21,23–26^ and likely also in monkey embryos ^25^.

In humans, XCI is established at peri-implantation stages with different dynamics depending on the lineage, occurring first in the trophectoderm ^20,23,27^. Similar findings have recently been reported for Cynomolgus monkey embryos ^25^. In addition, extraembryonic development differs markedly between mice and humans, both at the morphological and molecular levels, with humans having no extraembryonic ectoderm, in contrast to the mouse ^28^. Moreover, the extraembryonic mesoderm is specified after implantation and before gastrulation in humans, while in mice, these cells are only specified after gastrulation ^29–35^.

Addressing the key question of XCI mechanisms and requirements for human embryo development requires appropriate systems to model the establishment of XCI in humans ^36^. Naive human embryonic stem cells (hESCs) have emerged as an attractive system ^37,38^, as they represent the preimplantation human epiblast state, and carry two active X chromosomes (XaXa). Of note, in naive hESCs, *XIST* expression is typically observed from one of the two X chromosomes, in contrast to the pre-implantation embryo epiblast, where most female cells have two *XIST* clouds ^18,19^. *XIST* has recently been shown to trigger X-chromosome dampening in naive hESCs ^21,26,39,40^. *XIST*-coated X chromosomes are enriched in polycomb-mediated repressive histone modifications, histone H2A lysine 119 monoubiquitination (H2AK119ub) and histone H3 lysine 27 trimethylation (H3K27me3) in naive hESCs, as in embryos, despite their active states ^26^. The role of *XIST* in the establishment of XCI in humans is currently assumed and remains to be formally tested.

Naive hESCs can be differentiated toward the postimplantation epiblast lineage, a process termed capacitation ^41^. During this transition, cells acquire competence for somatic cell differentiation and eventually reach a primed pluripotent stem cell state, analogous to the post-implantation epiblast ^41^. Unlike their mouse counterparts, naive hESCs can also be differentiated into extraembryonic lineages such as postimplantation extraembryonic mesoderm cells (EXMCs) and trophoblast stem cells (TSCs), ^42–46^ in addition to all three founding lineages of the embryo^47–50^. Several lines of evidence indicate that XCI occurs in the course of *in vitro* differentiation of naive hESCs toward TSCs, trophoblast organoids and during capacitation ^51–53^. However, the functional and mechanistic interplay between XCI and extraembryonic fate induction has not been elucidated.

Here, we capitalized on *in vitro* cell fate induction to study the mechanisms and significance of XCI in early human development. We show that while one X chromosome acquires multiple hallmarks of XCI in *in vitro* derived extraembryonic cells, including gene silencing, exclusion of RNA polymerase II and a compact *XIST* RNA configuration, it adopts an unusual chromatin state that differs from that of embryonic cells, with limited enrichment in heterochromatin hallmarks compared to the active X chromosome and to autosomes. We further reveal that the *de novo* establishment of human XCI is closely linked, in time, with the emergence of EXMCs and TSCs. More strikingly, we show, using an inducible loss of function approach, that XCI is strictly dependent on *XIST* expression and that *XIST* impacts female extraembryonic cell survival. In addition to firmly establishing that *XIST* is necessary for the establishment of XCI in humans, our work provides strong evidence of XCI contribution to female human extraembryonic cell development.

## RESULTS

### XCI is established during the differentiation of naive hESCs toward extraembryonic lineages

To investigate and compare X-chromosome states in naive hPSCs and extraembryonic cell types, we differentiated female H9 naive hESCs into EXMCs and TSCs using trophoblast stem cell medium ASECRiAV or ACE (**Figure 1A**). After 30 days, we sorted distinct populations of EXMCs and TSCs using embryo extraembryonic mesoderm (EXM) and trophoblast markers BST2 and ENPEP, respectively (**Figure S1A**) ^44,47^. While the relative proportion of EXMCs and TSCs varied, naive hESCs and flow-sorted EXMCs and TSCs consistently displayed the expected dome-shaped, mesenchymal, and cuboidal morphologies, respectively **(Figure S1B).** We confirmed their identities using immunofluorescence (IF) for lineage markers (**Figure S1C**) and bulk RNA sequencing (RNA-seq) (**Figure S1D**), consistent with our previous study, in which only lineage-specific markers were used to distinguish TSCs and EXMCs while excluding other lineages ^44^.

**Figure 1.**
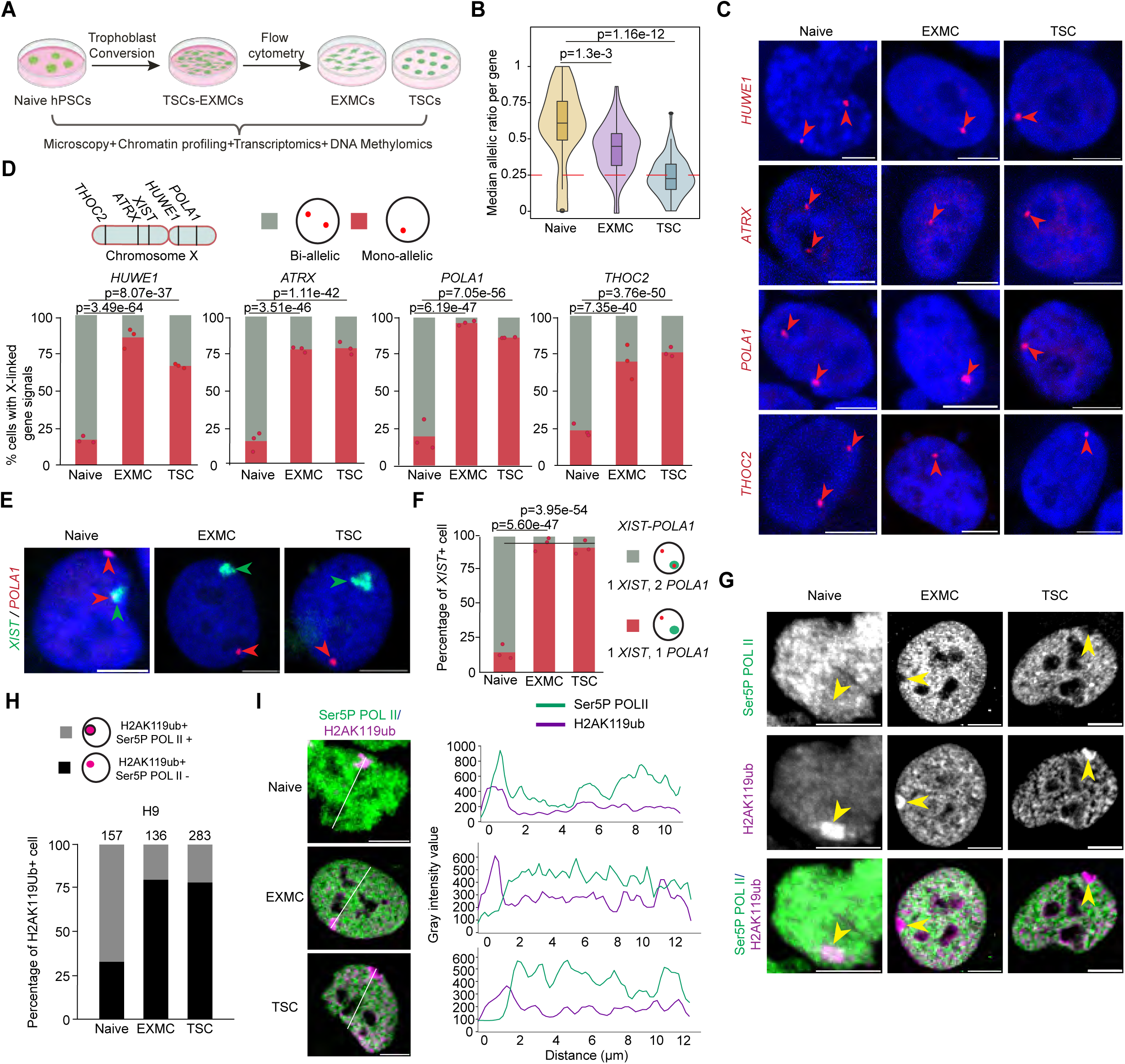
XCI is established during the differentiation of naive hESCs toward extraembryonic lineages. **A.** Schematic illustration showing naive hESCs to EXMCs and TSCs differentiation and assaying X-chromosome inactivation using several approaches. **B.** Violin plot showing average allelic ratio of X-linked genes in H9 naive hESCs (Passage 24), EXMC (Passage 6) and TSC (Passage 8) bulk RNA-seq data. There are 27, 35 and 32 informative genes in naive hESCs, EXMCs and TSCs respectively. Statistical significance was assessed using a t-test for comparison between naive vs EXMC and TSCs. P-values are shown in the figure. A p-value <0.05 was considered statistically significant. **C.** Representative RNA FISH images of four selected X-linked genes (Red) *HUWE1*, *ATRX*, *POLA1*, and *THOC2* in H9 naive (Passage 26) hESCs, EXMCs (Passage 6) and TSCs (Passage 9). Scale bar, 5 µm. **D.** Bar plots showing the quantification of the proportion of cells with monoallelic (Red) and biallelic (Gray) X-linked gene expression in H9 Naive, EXMC, and TSC. Data represents the mean from 3 independent RNA FISH experiments (n = 3). Each experiment was performed in triplicate. Statistical significance was assessed using a logistic regression test for comparisons between naive vs EXMC and naive vs TSC for each gene. P-values are shown in the figure. A p-value <0.05 was considered statistically significant. **E.** Representative RNA FISH images of *XIST* (Green) and *POLA1* (Magenta) in H9 naive (Passage 26), EXMC (Passage 6) and TSC (Passage 9). Scale bar, 5 µm. **F.** Bar plots showing the quantification of the proportion of cells with POLA1 puncta and *XIST* clouds in *XIST*-positive H9 Naive, EXMC, and TSC of the previous panel. Data represents the mean from 3 independent experiments (n = 3). Each experiment was performed in triplicate. Statistical significance was assessed using a logistic regression test for comparisons between naive vs EXMC and naive vs TSC. P-values are shown in the figure. A p-value <0.05 was considered statistically significant. **G.** Representative IF of Ser5P POL II (Green) and H2AK119ub (Magenta) in naive hESCs (Passage 24), EXMCs (Passage 6) and TSCs (Passage 9). Scale bar, 5 µm. Arrowheads indicate the H2AK119ub enriched X chromosome. **H.** Bar plots showing the quantification of the number of H2AK119ub with or without Ser5P POL II in H9 naive hESCs, EXMCs, and TSCs.The number of nuclei counted is indicated above each bar. **I.** Left panel: Representative IF images of Ser5P POL II (Green) and H2AK119ub (Magenta) in H9 naive hESCs, EXMCs and TSCs. Scale bar, 5 µm. Right panel: Representative line plots of single nuclei showing the gray intensity value of Ser5P POL II across a line passing through the H2AK119ub cloud. For panels D, F, and H, the number of nuclei counted is indicated in supplemental table S1.

To determine if XCI is induced during extraembryonic fate induction, we analysed bulk RNA-seq data with allelic resolution. We calculated the median allelic ratio per X-linked gene by dividing the least expressed allele by the most expressed one. As such, equal expression from the two X chromosomes should result in an allelic ratio of one, and we set the threshold for biallelic expression to 0.25, as previously established ^26^. Confirming the presence of two active X chromosomes, most X-linked genes were expressed biallelically in naive hPSCs (**Figure 1B**) ^24^. The average allelic ratio of the X chromosome was slightly lower than one and than autosomes (**Figures 1B and S1E**), due to *XIST*-dependent dampening of one X chromosome ^21,26^. The median allelic ratio was below 0.25 in TSCs, consistent with a switch from biallelic to monoallelic expression of X-linked genes, and thus XCI (**Figure 1B**) ^51^. In EXMCs, the allelic ratio was above that of TSCs, yet significantly lower than in naive hESCs (**Figure 1B**). We also found that the overall X chromosome expression levels were reduced in EXMCs and TSCs when compared to naive hPSCs, while the expression of chromosome 7, which has similar size and gene density, remained constant (**Figure S1F**). Together, these results suggest that induction of EXMC and TSC identities is accompanied by changes in X chromosome expression allelism and levels that are coherent with the establishment of XCI.

To further probe the allelic expression of X-linked genes at the single-cell level during naive to extraembryonic fate induction, we used RNA fluorescent *in situ* hybridisation (RNA FISH). We examined the expression of four X-linked genes distributed along the X chromosome, *HUWE1*, *ATRX*, *POLA1,* and *THOC2,* and observed a switch from biallelic expression in the naive state to monoallelic expression in both extraembryonic cell types, in agreement with one X chromosome being inactivated (n = 3 independent FISH experiments; logistic regression was used for statistical analysis; p-values are included in the figures) (**Figures 1C, 1D and Table S1**). Simultaneous RNA FISH analysis of *XIST* and *POLA1* confirms *POLA1* expression from the active X chromosome, which is not coated by *XIST* (**Figures 1E, 1F, and Table S1**). Similar results were obtained with three additional hPSC lines, including one hESC and two human induced pluripotent stem cell (hiPSC) lines (ICSIG-1 hiPSCs, SIG-1 hiPSCs, and WIBR2-MGT hESCs), grown in two different naive PSC culture media (**Figures S1G and S1H**). Moreover, we quantified the RNA FISH signal intensity of *POLA1*, comparing *XIST*-coated and non-coated X chromosomes in naive cells. We found that *POLA1* expression from the *XIST*-coated chromosome was significantly lower than from the non-coated allele, further supporting the occurrence of X-chromosome dampening in naive human pluripotent stem cells, in line with previous studies (**Figures S1I and S1J**) ^21,24,26^.

The inactive state of the *XIST*-coated X chromosome can be monitored at the single cell level by assessing the exclusion of the elongating form of RNA Polymerase II, phosphorylated on Serine 5 (Ser5P POL II) ^3^. We thus combined IF for Ser5P POL II with H2AK119ub or H3K27me3, to mark the *XIST*-coated X chromosome ^24,26^. We detected a single focal enrichment of H2AK119ub and H3K27me3 in around 30-75% of cells, irrespective of their cell state (**Figures 1G, S1K-S1N, S1O-S1Q**) ^24,26^. In naive hESCs, the Ser5P POL II signal was detected within the zone of H2AK119ub and H3K27me3 enrichment, in line with the active status of the X chromosome in the naive pluripotent stem cell state (**Figures 1G-1I and S1L-S1Q**). In contrast, in extraembryonic cells, Ser5P POL II was excluded from the H2AK119ub/H3K27me3 domain, suggesting the formation of a repressive compartment on the Xi, in which elongating RNA POL II is excluded (**Figures 1G, 1H, S1L-S1Q**). These results were confirmed by quantifying the intensity value of Ser5P POL II in the vicinity of H2AK119ub both in naive and extraembryonic cells (20 random nuclei per cell type were used for the quantification) (**Figures 1I, S1Q-S1R**).

Taken together, these results indicate that the transition of naive hESCs to extraembryonic lineages *in vitro* leads to XCI establishment, marked by the formation of a transcriptionally repressed compartment and X-linked gene silencing.

### Unusual chromatin remodeling accompanies XCI during extraembryonic specification

The results reported above suggest that the inactive X chromosome is marked by polycomb repressive complex (PRC)-repressive histone marks. The combination of *XIST* RNA FISH with IF for H3K27me3 and H2AK119ub indeed showed that in both naive hESCs and extraembryonic cells, the *XIST*-coated X chromosome is consistently marked by these repressive histone modifications (**Figures S2A and S2B**).

To explore chromosome-wide the distribution of repressive hPTMs that are typically enriched on the Xi, including H3K27me3, H2AK119ub and H3K9me3, we performed CUT&RUN in EXMCs and TSCs and compared to our previously published data of naive hESCs^26^. H3K27me3 and H2AK119ub enrichment profiles on the X chromosome were correlated across the three cell types, and more strongly between EXMCs and TSCs, but not with those of H3K9me3 (**Figure S2C**). Visualization of the profiles confirmed that hPTMs H3K27me3 and H2AK119Ub display similar distribution patterns on the X chromosome across all cell types (**Figure 2A**). Examination of hPTM enrichment of key developmental regulators, including *NANOG*, *TFAP2L1*, *CDX2*, *HAND1*, *GATA4*, and *HOXD1,* confirmed the expected H3K27me3 and H2AK119ub enrichment correlated with gene repression as expected and confirmed the validity of the data sets (**Figure S2D**).

**Figure 2:**
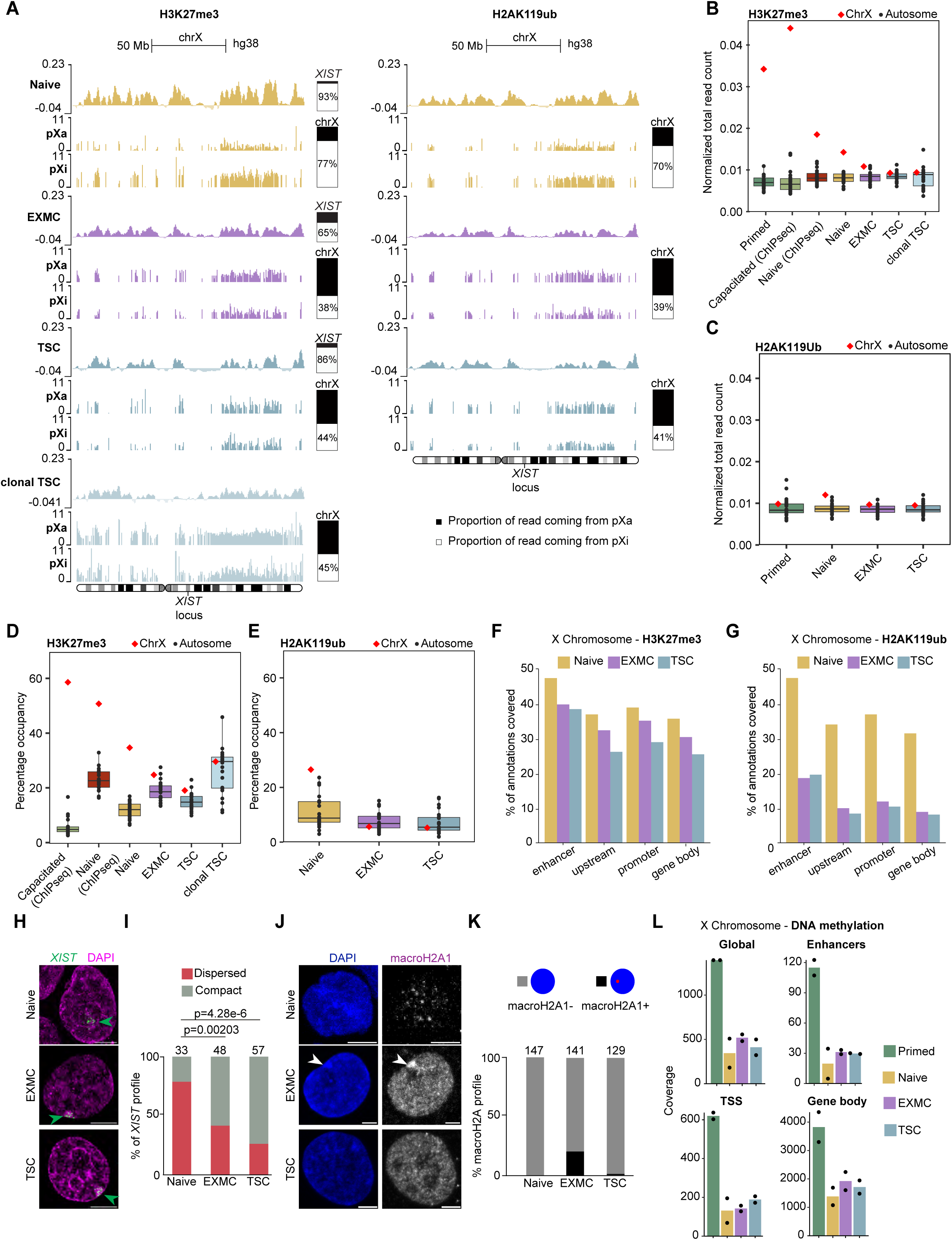
Unusual chromatin remodeling accompanies XCI during extraembryonic specification. **A.** CUT&RUN profiles (log2 enrichment over IgG) for H3K27me3 and H2AK119ub on the X chromosome in H9 naive hESCs (Passage 19), EXMCs (Passage 10) and TSCs (Passage 8). Allelic repartition of the informative reads are presented below each profile as reads mapped to the primed hESC active X (pXa) and the inactive X (pXi)^26^. The CUT&RUN experiments were performed in duplicate (n=2 replicates) on a WT clone. On the right of the profiles are depicted the allelic expression of *XIST* (top) and the allelic read quantification. **B.** Total number of CUT&RUN or ChIP-seq H3K27me3 reads from each chromosome, normalized by length of each chromosome in Primed hESCs ^26^, Capacitated hESCs ^26,53^, naive hESCs ^26,53^, H9 naive hESCs ^26^, EXMCs, TSCs and clonal TSCs. **C.** Total number of CUT&RUN H2AK119ub reads from each chromosome, normalized by the length of each chromosome in Primed hESCs ^26^, H9 naive hESCs ^26^, EXMCs and TSCs. **D.** Percentage of H3K27me3 CUT&RUN peak occupancy per chromosome in capacitated hESCs (ChIPseq data), naive hESCs (ChIPseq data), naive hESCs, EXMCs, TSCs and clonal TSCs. **E.** Percentage of H2AK119ub CUT&RUN peak occupancy per chromosome in H9 naive hESCs, EXMC and TSCs. **F.** Proportion (%) of annotation covered by CUT&RUN H3K27me3 peaks in H9 naive hESCs, EXMC and TSCs. **G.** Proportion (%) of annotation covered by CUT&RUN H2AK119ub peaks in H9 naive hESCs, EXMC and TSCs. **H.** Representative 3D-SIM images of *XIST* (Green) FISH and DAPI (Magenta) in H9 naive hESCs (passage 26), EXMCs (Passage 10) and TSCs (Passage 13). Scale bar, 5 µm. **I.** Bar plots showing the quantification of the proportion of cells with either compact or dispersed *XIST* clouds in H9 naive hESCs, EXMC, and TSC *XIST* RNA FISH, as shown in panel I. The number of nuclei counted is indicated below each bar. Statistical significance was assessed using a Chi^2^ test. P-values are shown in the figure. A p-value <0.05 was considered statistically significant. For details on how nuclei were classified as showing compact or dispersed signals, please refer to the Materials and Methods section. **J.** Representative IF of macroH2A1 in H9 naive hESCs (Passage 24), EXMCs (Passage 8) and TSCs (Passage 18). Scale bar, 5 µm. The experiment was performed only in one WT hESCs line (n=1). **K.** Bar plots showing the quantification of the proportion of macroH2A1 clouds in H9 naive hESCs, EXMCs and TSCs. Percentage of total cell count are shown. The number of counted nuclei are indicated above each bar. **L.** CpG islands methylation score on the X chromosome of H9 naive hESCs (Passage 28), EXMCs (Passage 7), TSCs (Passage 9) and primed hESCs (Passage 41) on the total X chromosome (top left), enhancers of the X chromosome (top right), transcription start sites of the X chromosome (TSS) (bottom left), and gene body of the X chromosome (bottom right). Number of replicates = 2 (n= 2).

Despite the shared pattern, normalized enrichment and normalized read distribution for H3K27me3 and H2AK119ub on the X chromosome were lower in EXMCs and TSCs compared to naive hESCs (**Figures 2A-2C**). To further understand chromatin changes upon XCI, we proceeded with allelic analysis of the CUT&RUN data. We used *XIST* RAP-seq data on clonal primed cells (with 100% skewed XCI^26,54^) and H9 whole genome sequencing data to generate a partly phased X-chromosome haplotype (12,813 X-linked SNPs), in which SNPs present in the *XIST* RAP-seq data were assigned to the Xi in the primed state (pXi). We first confirmed that both X chromosomes are equally accessible by CUT&RUN, as evidenced by equal representation of both X chromosomes in IgG control data (**Figure S2E**). In chemically reset naive hESCs, we observed a skewed enrichment of H3K27me3 and H2AK119ub marks (77% and 70%, respectively, of the X-linked reads) on the pXi, in agreement with XIST being expressed solely from this chromosome ^26^. In TSCs in contrast, despite a similar skewing of *XIST* expression (**Figure 2A**), these PRC-associated hPTMs were equally distributed on the two X chromosomes. Similar results were obtained on a clonal TSC sample, derived by differentiation of naive hESCs, followed by single TSC sorting and clonal TSC line expansion (**Figures S2F, S2G and S3A**).

To quantify the enrichment of hPTMs further, we counted the total number of reads for each condition, including in our analysis ChIP-seq datasets from naive and capacitated hESCs ^26,53^. In all naive hESCs samples, H3K27me3 and H2AK119ub were more enriched on the X chromosome compared to autosomes (**Figures 2B and 2C**), as previously reported^26^. Such enrichment of H3K27me3 was even more pronounced in primed and capacitated hESCs, likely reflecting lower levels of this mark on autosomes in primed hPSCs (**Figure 2B**)^55^. In contrast, in EXMCs and TSCs, H3K27me3 levels were of similar ranges on the X chromosome and autosomes (**Figures 2B and 2C**). Similar observations were made when independently generated H3K27me3 CUT&RUN or CUT&Tag data from H9-derived TSCs from another laboratory (H9 TSC II), embryo-derived TSCs (CT27)^42^, and clonal TSCs were analyzed (**Figures 2D, 2E, S3B-S3D**). We finally assessed whether more subtle changes in H3K27me3 and H2AK119ub patterns accompany XCI. We found that while the coverage levels of these marks across the genomic features studied (enhancers, gene upstream regions, promoters, and gene bodies) were lower in EXMCs and TSCs compared to naive hESCs, their relative distribution was similar in all cell types (**Figures 2F and 2G**). We then examined specifically X-linked genes that are monoallelically expressed and thus subjected to XCI in EXMCs and TSCs. In naive hESCs, EXMCs and TSCs, almost no enrichment in H3K27me3 and H2AK119ub could be seen on the gene bodies of *ATRX*, *POLA1,* and *THOC2*, in contrast with capacitated cells, whereas flanking regions tend to display higher enrichment in naive hESCs compared to EXMCs and TSCs (**Figures S3E and S3F**).

Collectively, these results suggest that the inactive, *XIST*-coated X chromosome is not particularly enriched in H3K27me3 (and H2AK119ub) compared to the other X chromosome and to autosomes in TSCs and EXMCs, in contrast to capacitated and primed pluripotent stem cells.

To investigate the apparent contradiction between IF and CUT&RUN results, we first asked whether the visible enrichment of repressive marks by IF was due to increased chromatin condensation. Quantifying DAPI intensity inside versus outside the *XIST*-coated X chromosome showed significantly higher condensation in extraembryonic cells compared to naive hESCs (**Figures S4A–S4C**). In addition, exploring the global increase of repressive marks genome-wide revealed that H3K27me3 IF intensity was elevated across the nucleus in extraembryonic cells relative to naive hESCs (**Figures S4D–S4E**). Together, these findings suggest that the accumulation of PRC-mediated repressive marks on the Xi observed by IF is due to X chromosome compaction rather than relative enrichment compared to other chromosomes.

This is in line with the pattern of *XIST* expression. Indeed, differences in *XIST* distribution between naive and post-XCI states have been reported^20^. In human pre-XCI cells (preimplantation embryos and naive hESCs), *XIST* RNA adopts a dispersed configuration, while embryonic post-XCI cells such as primed pluripotent stem cells and fibroblasts show a compact *XIST* RNA cloud^20,21^. To probe *XIST* spreading and compaction, we employed RNA FISH followed by both conventional confocal microscopy and super-resolution three-dimensional structured illumination microscopy (3D-SIM) and measured the volume and diameter of the *XIST* RNA FISH signal in naive hESCs, EXMCs and TSCs. A single *XIST* cloud was detected in a vast majority of naive hESCs, as previously reported ^21,24,26,54^, as well as in EXMCs and TSCs (**Figures S4F-S4I**). We found that the transition of naive cells to extraembryonic cell fates were associated with the formation of more compact *XIST* clouds, as in post-XCI cells of embryonic origin (**Figures 2H-2I, S4J-S4K, Videos S1, S2, and S3**). Moreover, using 3D-SIM imaging, we quantified the number of *XIST* foci in naive hESCs, EXMCs, and TSCs. Naive cells exhibited approximately 50 foci per nucleus, while EXMCs and TSCs showed around 30 foci per nucleus (**Figure S4L**).

To further characterize the chromatin state of the Xi in EXMCs and TSCs, we investigated other chromatin features. We examined, using IF, the presence of macroH2A, a histone variant recruited in the late stages of XCI during differentiation of mouse embryonic stem cells^56–59^, and previously shown to mark the Xi in primed hESCs^60,61^. Only a minority of EXMCs and TSCs displayed macroH2A enrichment on the Xi (1% and 20%, respectively) (**Figures 2J-2K**). We also assessed CpG island methylation, a mark commonly associated with promoter regions of the Xi primed hESCs and somatic cells^60–63^, using methylated DNA sequencing (MeD-seq)^64^. We found that the DNA methylation levels on the X chromosome of EXMCs and TSCs were much lower than in primed hESCs and comparable to those of naive hESCs (**Figure 2L)**. These results suggest that the X chromosome is hypomethylated in EXMCs and TSCs compared to the Xi of embryonic lineage cells.

Collectively, these results showed that, in human extraembryonic cells, the Xi displays an atypical chromatin state, with the absence of most chromatin hallmarks of XCI.

### The establishment of XCI is closely linked with the acquisition of extraembryonic identity

To link in time extraembryonic induction and the establishment of XCI, we first analyzed the kinetics of XCI during the transition from naive hESCs to EXMCs and TSC. Samples were collected on days 0, 4, 8, 10, 12, 14, and 16 of differentiation (**Figure 3A**). RNA FISH for the X-linked gene *POLA1*, used as a first proxy for XCI, revealed a pronounced transition from biallelic to monoallelic expression between days 4 and 8 (data represent the mean from 3 independent time-course differentiation experiments followed by IF-FISH) (**Figures 3A, 3B, and Table S1**). A similar trend was obtained with another X-linked gene, *HUWE1*, analyzed in H9 cells resetted and maintained in different naive culture conditions (**Figures S5A-S5B**). We next integrated, by combining IF and RNA FISH, allelic analysis of *POLA1* expression with expression of trophoblast and embryo EXM markers, GATA3 and VIM, respectively ^44^. GATA3+ and VIM+ cells were first detected at day 8 (**Figures 3C-3F**), as previously reported ^44^. *POLA1* was monoallelically expressed in the majority of EXMCs and TSCs from day 8 onwards (**Figure 3B**).

**Figure 3:**
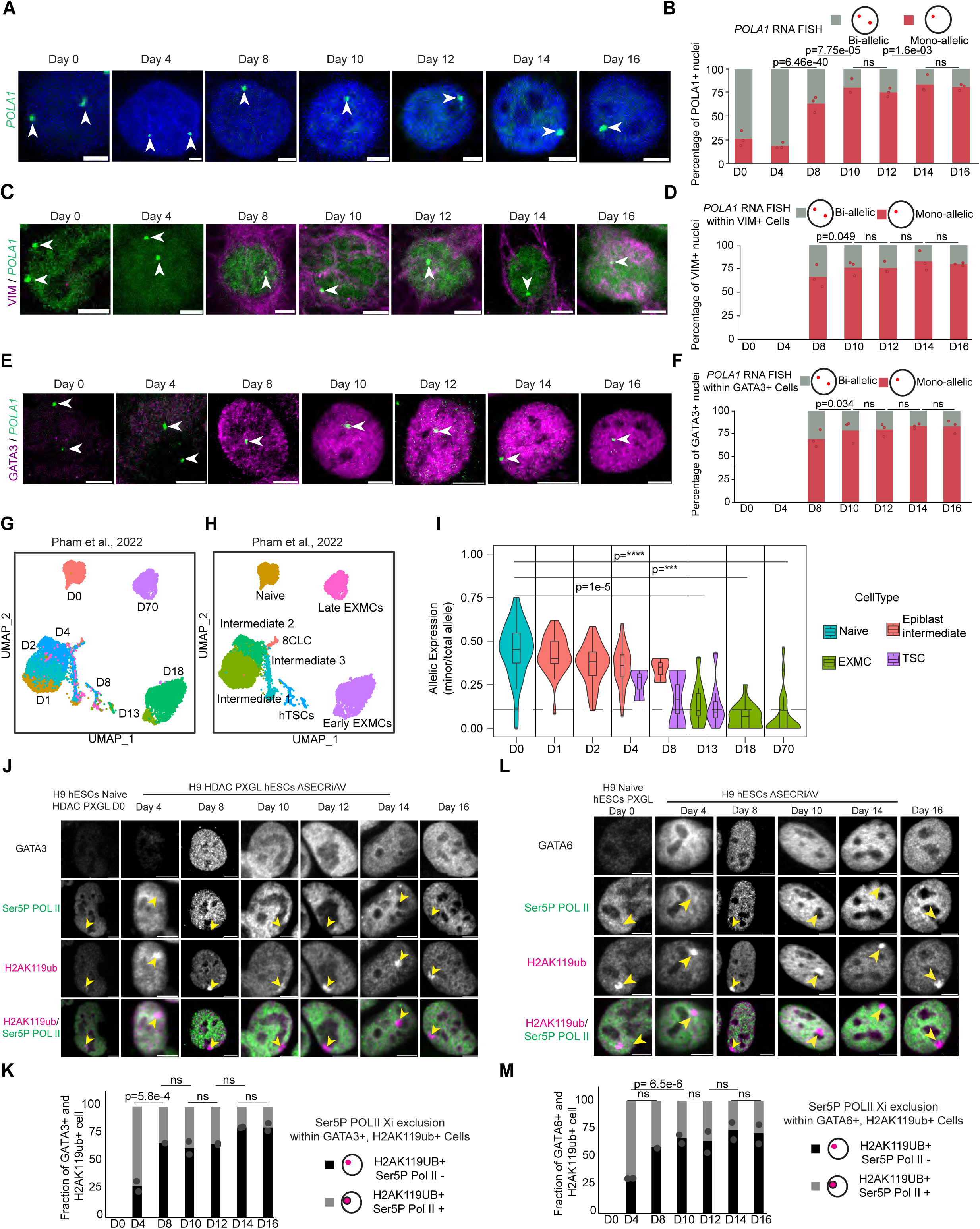
The establishment of XCI is closely linked with the acquisition of extraembryonic identity. **A.** Representative X-linked gene FISH images showing *POLA1* puncta dynamic during edification of H9 naive hESCs (Passage 27) to EXMCs and TSCs. Samples were collected at seven different time points, as indicated. Scale bar, 5 µm. **B.** Bar plots showing the quantification of the proportion of cells with *POLA1* puncta during the differentiation of H9 naive hESCs to EXMCs and TSCs in the previous panel. The number of nuclei counted is indicated in supplemental table S1 for each round of experiments. Data represents the mean from 3 independent time-course differentiation experiments (n= 3) followed by three independent FISH experiments (n= 3). Each experiment was performed in triplicate. Statistical significance was assessed using a logistic regression test for comparisons between different time points. P-values are shown in the figure. A p-value <0.05 was considered statistically significant. **C.** Representative IF-FISH of X-linked gene *POLA1* (Green) and EXMC lineage marker VIM (Magenta) images showing dynamics during H9 naive hESCs to EXMCs and TSCs differentiation. Samples were collected at seven different time points, as indicated. Scale bar, 5 µm. **D.** Bar plots showing the quantification of the proportion of cells with *POLA1* puncta during the differentiation of naive hESCs to H9 EXMCs and TSCs in the previous panel within the VIM positive nuclei. The number of nuclei counted is indicated in supplemental table S1 for each round of experiments. Data represents the mean from 3 independent time-course differentiation experiments (n= 3) followed by three independent IF-FISH experiments (n= 3). Each experiment was performed in triplicate. Statistical significance was assessed using a logistic regression test for comparisons between different time points. P-values are shown in the figure. A p-value <0.05 was considered statistically significant. **E.** Representative IF-FISH of X-linked gene *POLA1* (Green) and trophoblast lineage marker GATA3 (Magenta) images showing dynamics during H9 naive hESCs to EXMCs and TSCs differentiation. Samples were collected at seven different time points, as indicated. Scale bar, 5 µm. **F.** Bar plots showing the quantification of the proportion of cells with *POLA1* puncta during the differentiation of naive hESCs to H9 EXMCs and TSCs in the previous panel within the GATA3 positive nuclei. The number of nuclei counted is indicated in supplemental table S1 for each round of experiments. Data represents the mean from 3 independent time-course differentiation experiments (n= 3) followed by three independent IF-FISH experiments (n= 3). Each experiment was performed in triplicate. Statistical significance was assessed using a logistic regression test for comparisons between different time points. P-values are shown in the figure. A p-value <0.05 was considered statistically significant. **G.** UMAP of time course scRNA-seq reanalysis of ^44^ data during differentiation of ICSIG-1 naive hiPSCs to EXMCs and TSCs as well as day 70 EXMCs, colored by days. **H.** As in 3G but colored by cell types. **I.** Plot showing allelic expression from a timecourse scRNA-seq dataset, calculated using the scLinaX pipeline ^65^. Only genes that we expect to be subject to XCI are included, and only cell types with at least 2 informative SNPs in a day are shown. Reanalysis of data from ^44^. Statistics calculated using a students t-test. *** tests were significant beyond the limits (<e^-16^) of the test to calculate. **J.** Representative IF Ser5P POL II (Green), H2AK119ub (Magenta), and trophoblast lineage marker GATA3 (Gray) images showing dynamics during naive to TSCs and EXMCs differentiation. Samples were collected at six different time points, as indicated. Scale bar, 5 µm. **K.** Representative bar plot showing the quantification (%) of the number of Ser5P POL II enriched nuclei during the differentiation of naive to TSC and EXMCs within the GATA3 and H2AK119ub double positive nuclei. Statistical significance was assessed using a logistic regression test. P-values are shown in the figure. A p-value <0.05 was considered statistically significant. **L.** Representative IF Ser5P POL II (Green), H2AK119ub (Magenta), and EXM lineage marker GATA6 (Gray) images showing dynamics during naive to TSCs-EXMCs conversion. Samples were collected at six different time points, as indicated. Scale bar, 5 µm. **M.** Representative bar plot showing the quantification (%) of the number of Ser5P POL II enriched nuclei during the differentiation of naive to TSC and EXMCs within the GATA6 and H2AK119ub double positive nuclei. Statistical significance was assessed using a logistic regression test. P-values are shown in the figure. A p-value <0.05 was considered statistically significant. For panels B, D, F, K, and M, the number of nuclei counted is indicated in supplemental table S1.

We also took advantage of our previously published single-cell RNA sequencing (scRNA-seq) time course data of naive hESCs to EXMCs and TSCs induction ^44^ to narrow down the relationship between X-linked gene expression and cell identity. We calculated the X-to-autosome ratio as a readout for XCI. This ratio remained constant up to day 4 of differentiation, then dropped significantly from day 8 onwards in cells identified as EXMCs and TSCs (**Figures 3G–3H and S5C–S5E).**

To further define the kinetics of XCI, we performed allelic analysis of scRNA-seq data using a recently developed bioinformatics tool, single-cell level inactivated X-chromosome mapping (scLinaX) ^65^. We observed a significant decrease of the allelic ratio for genes expected to be subject to XCI ^66^ from day 4 of differentiation (**Figure S5F**). By contrast, genes expected to escape XCI^66^ remained biallelically expressed throughout the time course (**Figures S5F-S5G**). When we further examined allelic X-linked gene expression in individual cell types, there was a significant decrease in EXMCs and TSCs by day 13 onwards, confirming the establishment of XCI (**Figure 3I**).

Time-course analysis of Ser5P POL II staining combined with that of GATA3 and GATA6 (as nuclear markers of TSCs and EXMCs, respectively) confirmed these results: the proportion of cells with Ser5P POL II exclusion from the H2AK119ub-marked chromosome increased most prominently between days 4 and 8 (**Figures S5H-S5I**). We further combined the above staining with immunostaining of GATA3 and GATA6 to detect the dynamics of TSCs and EXMCs, respectively, during the differentiation (**Figures 3J-3M**). Immunostaining analysis revealed that GATA3 and GATA6 positive cells started appearing between days 4 and 10, coinciding with the occurrence of Ser5P POL II exclusion, which suggests the formation of a repressive compartment (**Figures 3J-3M**). Collectively, these results suggest that X-chromosome silencing is closely linked with acquisition of extraembryonic cell identity, occurring concurrently with or shortly after differentiation.

### Loss of *XIST* dramatically impacts human extraembryonic cell survival

The correlation, in time, between the establishment of XCI and the emergence of extraembryonic cells prompted us to investigate the functional interconnection between these two processes. To do so, we exploited a conditional *XIST* KD system (*XIST* iKD), in which *XIST* repression can be triggered upon doxycycline (Dox) treatment by the recruitment of dCas9-KRAB (**Figures 4A and S6A**) ^26^. We confirmed that a ten-day treatment of naive hESCs with Dox (+Dox) resulted in nearly complete repression of *XIST* in three independent clones (**Figures S6B-S6D**). *XIST* repression did not affect the expression of naive pluripotency genes (**Figure S6D**), and naive hESCs lacking *XIST* expression proliferated similarly to wild-type (WT) cells (**Figure S6E**) ^21,26^. By contrast, *XIST* repression had a pronounced effect during the differentiation of naive hESCs toward EXMCs and TSCs. *XIST*-depleted and WT cells were indistinguishable during the initial steps of differentiation. However, most of the *XIST*-depleted cells were lost by day 15 of extraembryonic fate induction, leaving only feeder cells visible as identified by the presence of chromocenters (**Figures 4B-4C).** Similar results were obtained in multiple *XIST* iKD clones **(Figure S6F**). In the absence of *XIST*, no GATA3+ colonies could be observed at day 15, whereas GATA3+ cells were detectable in WT cells treated or not with Dox (**Figures 4D and S6G**). The striking loss of trophoblast cells upon *XIST* repression was confirmed using flow cytometry analysis for the trophoblast marker ENPEP at day 15 (**Figure 4E**). To establish if the observed phenotype was linked to the absence of *XIST* expression during the differentiation or was a consequence of *XIST* loss in naive hESCs, we included a condition where *XIST* was depleted in naive cells, followed by Dox removal from the media from day 0 of differentiation. This allowed for *XIST* re-expression, which partially rescued the phenotype, with around 18% of cells expressing ENPEP at day 15, compared to 35% in untreated conditions (**Figure S6H-S6I**).

**Figure 4:**
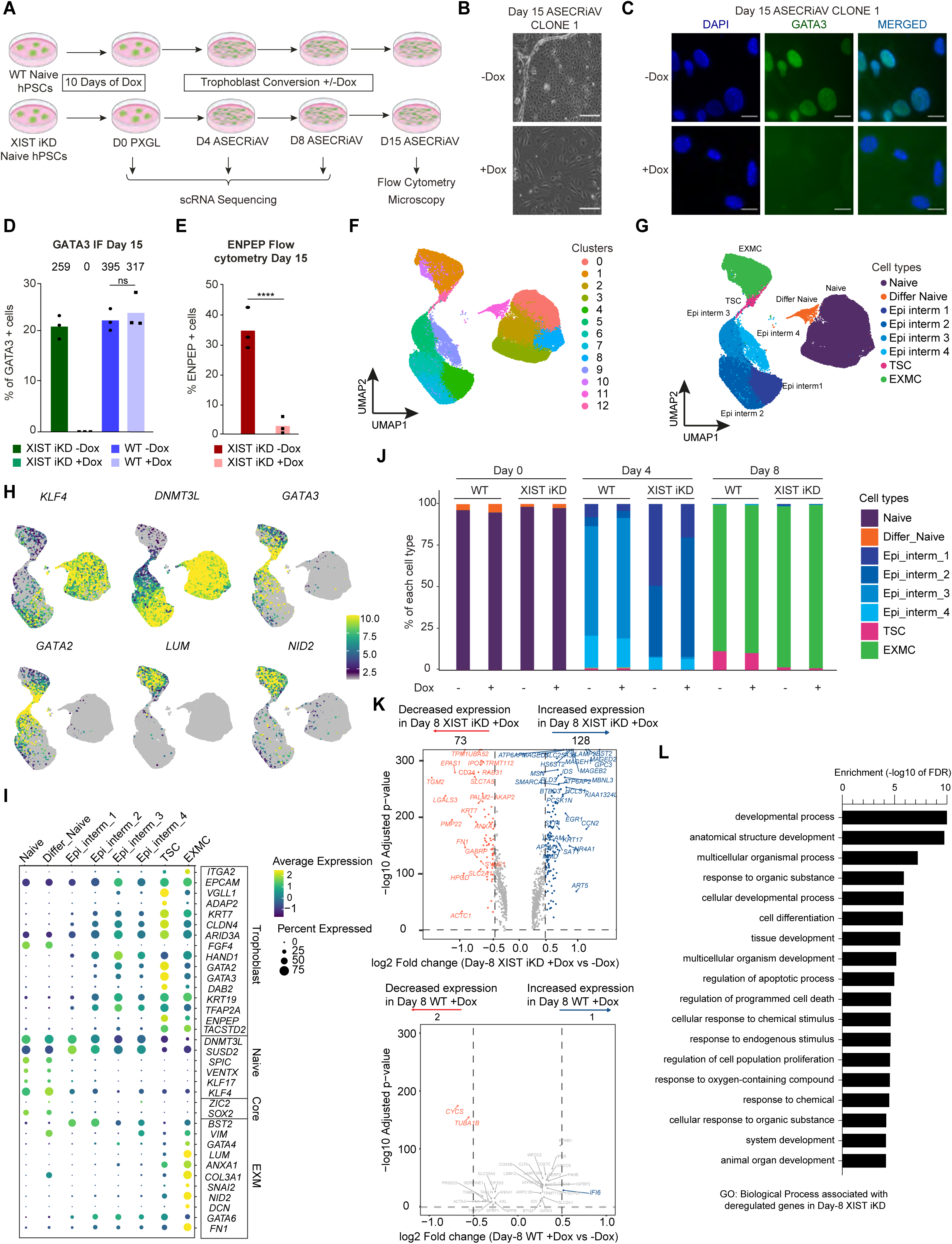
Loss of *XIST* dramatically impacts human extraembryonic cell survival. **A.** Schematic illustration showing wild type (WT) and *XIST* inducible knock down (iKD) H9 naive hESCs under trophoblast differentiation medium with or without Doxycycline (Dox) and assaying cell identity and X-chromosome inactivation using microscopy, scRNA-seq, bulk RNA-seq and flow cytometry. **B.** Representative bright-field microscopy images showing the morphology of H9 *XIST* iKD clone-1 hESCs at Day 15 of culture in trophoblast differentiation medium with or without Dox. Note: Feeders only are seen in the +Dox conditions. Scale bar, 200 µm. The experiment was performed in 3 independent *XIST* iKD clones of H9 hESCs. Out of which one is shown here and the other two are shown in figure S5E (total n=3 independent clones). **C.** Representative IF of GATA3 (Green) of cells mentioned in 4B. DAPI is shown in blue. Scale bar, 20 µm. The experiment was performed in 3 independent *XIST* iKD clones of H9 hESCs. Out of which one is shown here and the other two are shown in figure S5F (total n=3 independent clones). **D.** The proportion of GATA3 positive nuclei in both WT and *XIST* iKD H9 hESCs at Day 15 of trophoblast differentiation with or without Dox. The number of nuclei counted is indicated above each bar. Each dot represents one of three independent *XIST* iKD clones (n=3) or WT clones (n=3). Statistical significance was assessed using a t-test. A p-value <0.05 was considered statistically significant. The number of nuclei counted is indicated in supplemental table S1. **E.** Flow cytometry quantification of the proportion of ENPEP positive cells at day 15 of trophoblast differentiation of *XIST* iKD hESCs with or without Dox. Statistical significance was tested using a Chi² test, resulting in p<0,0001. Each dot represents one of three independent *XIST* iKD clones (n=3). **F.** UMAP of scRNA-seq time course during differentiation of H9 WT and *XIST* iKD hESCs with or without Dox under trophoblast culture conditions, 13 clusters are shown using different colors. The scRNA-seq data was generated using one of the above 3 clones both for the WT and the *XIST* iKD hESCs with or without Dox (n=1). **G.** As in 4F but colored by cell types. Lineage information where Naive = Naive epiblast, Differ Naive = Differentiated naive epiblast, Epi interim 1 = Epiblast intermediate type 1, Epi interim 2 = Epiblast intermediate type 2, Epi interim 3 = Epiblast intermediate type 3, Epi interim 4 = Epiblast intermediate type 4, TSC = Trohpblast stem cells, and EXMC = Extraembryonic mesoderm cells. **H.** UMAPs from Figure 4F showing the expression of *KLF4 and DNMT3L* (naive hESC markers), *GATA3 and GATA2* (TSC markers), *LUM* and *NID2* (EXMC markers) in time course scRNA-seq data. **I.** Dot plot showing the expression of marker genes for core pluripotency, naive pluripotency, trophoblast and embryo extraembryonic mesoderm (EXM) in time course scRNA-seq data from this study. **J.** The proportion of each cell type at different time points during differentiation of naive hESCs into TSCs and EXMCs. Colored by cell types. **K.** Differential gene expression as detected by scRNA-seq between Day 8 *XIST* iKD (Top) and WT cells (Bottom) with Dox vs without Dox. Dashed lines indicate -log10 adjusted p-Value <0.05 and log2 fold change < -0.5 or > 0.5. **L.** GO enrichment analysis of differentially expressed genes as a result of *XIST* depletion comparing Day 8 *XIST* iKD cells with and without Dox.

To gain deeper cellular and molecular insights into the origin of the phenotype resulting from *XIST* depletion, we conducted time course scRNA-seq analysis in *XIST* depletion and control conditions. We harvested cells at day 0, 4 and 8 of extraembryonic fate induction, before massive cell loss occurs in *XIST*-depleted conditions (**Figure 4A**). Uniform Manifold Approximation and Projection (UMAP) analysis of all conditions identified 13 clusters (**Figure 4F**). At day 0, the majority of cells expressed markers of naive pluripotency and thus corresponded to naive hESCs, irrespective of the conditions; a minimal fraction of cells displayed *VIM* expression together with lower expression of markers of naive pluripotency, which are categorized as differentiated naive cells, in line with a previous study (**Figures 4F-4J, S6J-S6K**) ^44^. By day 4, most cells appeared to progress into an epiblast intermediate state (**Figures 4F-4G)**, which is distinct from naive and primed epiblast states, as described previously ^44^. Epiblast intermediates display decreased expression of several naive pluripotency genes, including *KLF4, SPIC,* and *VENTX* while maintaining expression of formative pluripotency genes (**Figures 4H-4I, S6L and Table S2**). These epiblast intermediate populations formed four different clusters, which we termed epiblast intermediates 1, 2, 3, and 4. There was a slight variation in the proportion of each epiblast intermediate between the conditions (compare *XIST* iKD +/- Dox, **Figure 4J and Table S2**), which may reflect a role for *XIST* in the transition between distinct pluripotent epiblast states. On day 8, EXMCs and TSCs became the only cell types (**Figures 4G-4J, S6J-S6L and Table S2**). No differences were observed in the proportions of EXMCs and TSCs between Dox treated and untreated conditions, both in WT and *XIST* iKD cells (**Figure 4J and Table S2**). Altogether, these results suggested that in the absence of *XIST* expression naive hESCs can exit pluripotency and enter extraembryonic cell identities, marked by the acquisition of a transcriptional program characteristic of EXMCs and TSCs. However, upon passage, these cells did not survive.

To infer the gene network impacted by *XIST* loss, we identified differentially expressed genes (DEGs) between *XIST*-positive and *XIST*-negative cells at day 8 of differentiation, prior to *XIST*-depleted cell death. We found 73 genes with decreased expression and 128 genes with increased expression upon *XIST* depletion (**Figure 4K** and **Table S3**). In contrast, there were only three genes differentially expressed in WT Dox treated versus untreated cells, indicating that Dox treatment *per se* does not perturb gene expression (**Figure 4K** and **Table S3**). Gene ontology (GO) analysis revealed that deregulated genes in *XIST-*depleted cells are primarily implicated in development, cell differentiation, cell proliferation, and the regulation of apoptotic processes (**Figure 4L** and **Table S3**). These pathways are radically different from those corresponding to DEGs identified at day 0 and day 4, the latter instead concerning metabolic processes (**Figures S6M-S6O** and **Table S4**) and are in agreement with the phenotype observed.

Analysis of apoptosis-related gene expression confirmed global deregulation of this pathway upon *XIST* loss. Indeed, more than 30 of such genes display increased or decreased expression levels in *XIST* knockdown conditions (**Figures S7A-S7D**). Finally, we performed flow cytometry analysis at day 15 of ASECRiAV differentiation, combining ENPEP and Annexin V staining to mark the presence and relative proportion of TSCs and apoptotic cells, respectively (**Figures S7E-S7F**). As expected, very few ENPEP+ cells were recovered in *XIST* iKD +Dox conditions. The few remaining ENPEP+ cells, however, display higher Annexin V staining, a marker of apoptosis, compared to control conditions. (**Figures S7E-S7F**). Collectively, these results strongly suggest that extraembryonic cells enter into apoptosis in the absence of *XIST* expression.

### *XIST* is necessary for the establishment of human XCI

We next probed more specifically whether the establishment of XCI was impaired by the lack of *XIST* expression and could thus be causally linked to the defect in extraembryonic cell survival. We aggregated scRNA-seq data per condition to generate pseudo-bulk datasets and calculated the mean X chromosome-to-autosome (X:A) expression ratio. This ratio was strongly elevated upon *XIST* depletion compared to control conditions, in particular at day 8 of differentiation, where it was most pronounced. A mild increase was also observed at days 0 and 4, which might be due to the role of *XIST* in X-chromosome dampening^26^ (**Figure 5A**). Subcategorization of day 8 cells as EXMCs and TSCs showed a marked increase in X:A ratio upon *XIST* depletion in both cell types (**Figure 5B**).

**Figure 5:**
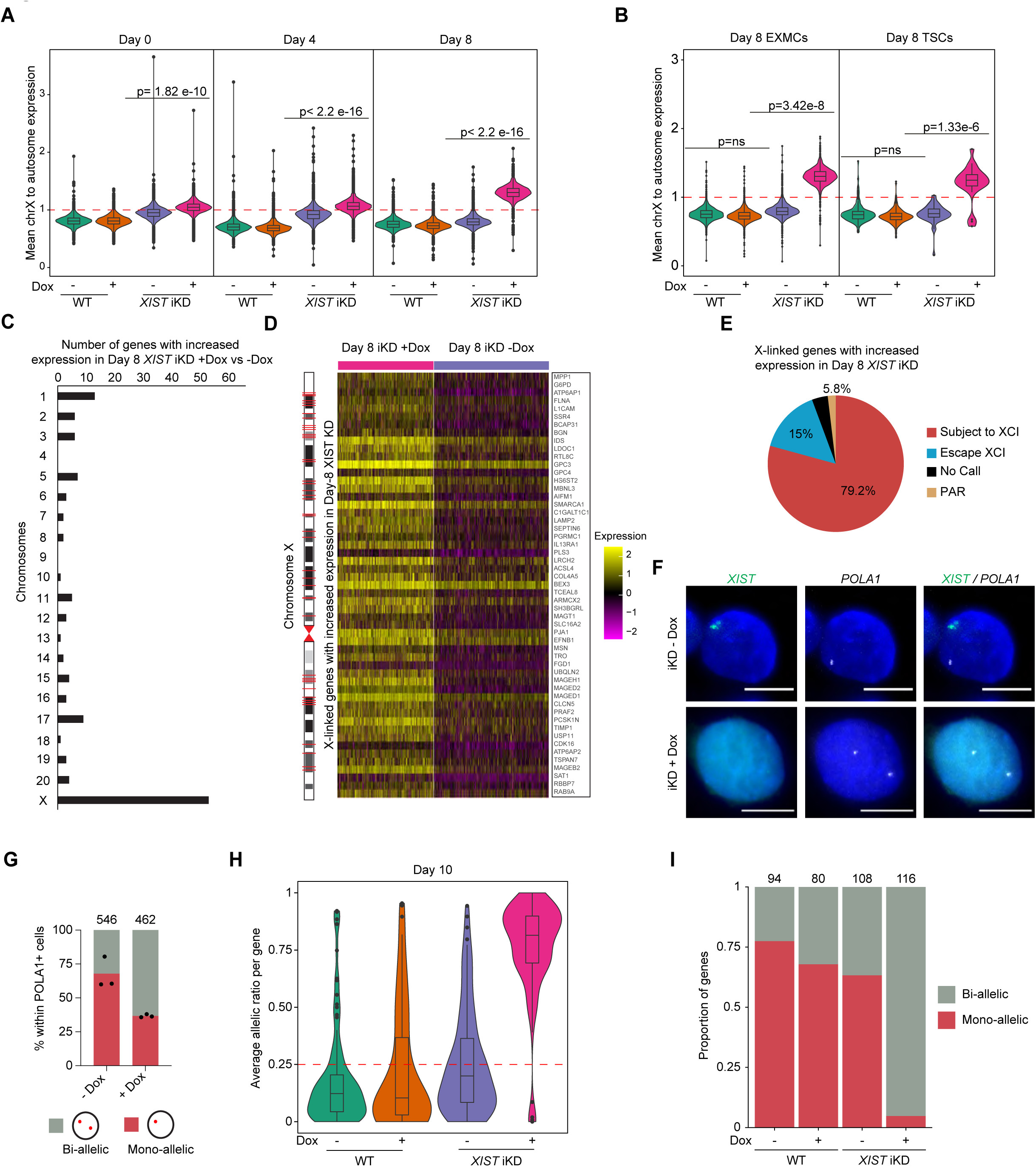
Human *XIST* is necessary for the establishment of human XCI in extraembryonic cells. **A.** Mean X-chromosome-to-autosome gene expression at different time points during ASECRiAV differentiation of WT and *XIST* iKD H9 hESCs with and without Dox. Colored by conditions. Statistical significance was evaluated using a t-test on a randomly downsampled dataset (100 cells per condition) to mitigate the influence of the large sample size, which otherwise yielded highly significant p-values across all comparisons. Bonferroni-corrected p-values are displayed in the figure, with p < 0.05 considered statistically significant. In cases where the p-value was exceedingly small, it is reported as p < 2.2e-16, reflecting the detection limit of the statistical software. **B.** Mean X-chromosome-to-autosome gene expression in H9 TSCs and EXMCs in Day 8 of WT and *XIST* iKD conditions with or without Dox. Statistical significance was evaluated using a t-test on a randomly downsampled dataset (24 cells per condition) to mitigate the influence of the large sample size, which otherwise yielded highly significant p-values across all comparisons. Bonferroni-corrected p-values are displayed in the figure, with p < 0.05 considered statistically significant. **C.** Number of DEGs on each chromosome in Day 8 *XIST* iKD without *XIST* (+Dox) versus with *XIST* (-Dox). **D.** The expression of all X-linked DEGs in day 8 *XIST* iKD H9 cells with and without *XIST* (Right panel). The X chromosome map view shows the relative location of the X-linked DEGs on the chromosome (left panel). **E.** Proportion of X-linked DEGs in Day 8 *XIST* depleted H9 hESCs belonging to different categories of X-linked genes based on their known XCI status in somatic cells_67._ **F.** Representative RNA FISH image of *XIST* and *POLA1* in Day 8 *XIST* iKD with Dox (*XIST*-) or without Dox (*XIST*+). Scale bar, 10µm. **G.** Proportion of biallelic and monoallelic *POLA1* signal on day 8 *XIST* iKD with Dox (*XIST*-) or without Dox (*XIST*+). The number of counted nuclei is indicated above each bar. The experiment was replicated on the three independent clones. The number of nuclei counted is indicated in supplemental table S1. **H.** Violin plot showing average allelic ratio per gene in Day 10 WT and *XIST* iKD with or without Dox, on a *XIST* iKD or a WT clone (n=1). **I.** Proportion of X-linked genes showing biallelic and monoallelic expression in Day 10 WT and *XIST* iKD cells with or without Dox. The number of genes analyzed is indicated above each bar.

To test whether the elevated X:A ratio in the absence of *XIST* resulted from downregulated X-linked genes, we mapped to the genome, among the DEGs identified at day 8 of differentiation, the genes displaying increased expression in *XIST* repressed conditions (hereby referred to as increased expression genes, IEGs). This revealed a strong enrichment of IEGs on the X chromosome (41.5%) relative to other chromosomes (**Figures 5C, S8A and Table S5**). This contrasts with the chromosomal distribution of IEGs identified at day 0 and day 4 of differentiation, of which only 5.6% and 11.8% were X-linked, respectively (**Figures S8B-S8C and Table S5**). The day 8 X-linked IEGs were distributed along the entire X chromosome (**Figure 5D and Table S5**). Moreover, most of these genes (79.2%) were defined as subjected to XCI in a comprehensive meta-analysis performed in somatic cells ^67^ (**Figure 5E**). Another 15% of IEGs were reported as escapees, which can be explained by a differential escaping pattern in embryonic versus extraembryonic lineages and/or by the influence of *XIST* on the expression levels of escapees, in line with a recent study in mice ^68^. Collectively, these results indicate that human *XIST* is required for the establishment of chromosome-wide X-linked gene silencing during extraembryonic cell fate induction.

To formally determine the effect of *XIST* depletion on XCI, we first assessed the expression of the *POLA1* X-linked gene at single-nucleus and allelic resolution by RNA FISH. The majority of day 8 *XIST-*depleted cells displayed biallelic *POLA1* expression, while this gene was silenced on the *XIST*-coated chromosome in control conditions (**Figures 5F-5G**). To probe, at the chromosome scale, the impairment in XCI establishment in the absence of *XIST*, we performed new bulk RNA-seq and allelic analyses. Gene expression analyses confirmed the induction of EXMC and TSC markers in all four samples at day 10 of differentiation, the last time point before cells died (**Figure S8D**). In all samples in which *XIST* is expressed (WT -Dox, WT +Dox and *XIST* iKD -Dox), the mean allelic ratio was below 0.25, indicating monoallelic X-linked gene expression, hence XCI (**Figure 5H**). This contrasted with *XIST* depleted samples, which displayed an allelic ratio of 0.8, reflecting biallelic gene expression, hence the lack of XCI. We then categorized genes as mono- or biallelically expressed using an allelic ratio threshold of 0.25. This confirmed that, in the presence of *XIST*, most X-linked genes were subjected to XCI at day 10, as inferred from their monoallelic expression pattern, and that this process is severely impaired in the absence of *XIST,* with most genes remaining biallelically expressed (**Figure 5I**). These results demonstrate that *XIST* is required for the establishment of XCI in humans. They also suggest that the phenotype we observed resulted from impaired XCI.

## DISCUSSION

Mice have been for long the model of choice for investigating XCI mechanisms and function, with the implicit assumption that these would stand for all mammals. XCI studies in non-rodent species have, however, challenged this premise and highlighted the need to revisit the regulation of X-chromosome dosage and its consequences in a species-specific manner. There are obvious interests in addressing these questions in humans, and there are also major limitations in terms of relevant biological resources and opportunities for functional studies in early developmental contexts.

The advent of stem cell biology and the emerging ability to derive and maintain naive hPSCs which recapitulate, *in vitro*, preimplantation stages of development paved the way for XCI studies in humans ^69^. Our study establishes naive hPSCs as a relevant and powerful system to dissect the mechanistic and functional aspects of *de novo* XCI establishment in humans. Here we exploited their unique property, the ability to give rise, *in vitro*, to post-implantation extraembryonic cells of different natures, TSCs and EXMCs, to link XCI and extraembryonic fate in humans. Our work opens avenues for understanding more generally the role of XCI in lineage induction and in human embryo development.

Our results reveal that induction of extraembryonic fates from naive hPSCs recapitulates primate-specific biology, such as the transition from X-chromosome dampening to XCI that takes place in early human embryogenesis ^18,23,27^. We indeed confirmed, using a combination of approaches that provide gene-specific as well as chromosome-wide assessment of silencing, that XCI is established in *in vitro* generated TSCs and EXMCs, in line with previous findings ^51,52^. A peculiar observation we made concerned the features of the Xi in extraembryonic cells, which is not enriched in chromatin repressive hallmarks as compared to the autosomes and to the Xa when assessed by CUT&RUN analysis, a conclusion that was confirmed in independent analyses of additional TSC lines. The DNA hypomethylation we report is also in line with X-linked gene promoter hypomethylation in human female placentas^61^. That the Xi was labeled with anti-H2AK119Ub and H3K27me3 antibodies in EXMCs and TSCs in IF experiments might be explained by the overall higher density of histones on this chromosome ^70^ and/or by the peculiar Xi 3D conformation ^71,72^. Indeed, we detected increased chromatin compaction of the Xi relative to autosomes in EXMCs and TSCs, as well as increased autosomal levels of H3K27me3 and H2AK119ub, which may explain the lack of detectable relative enrichment of these hPTMs by CUT&RUN analysis. It is intriguing to note that the Xi was also reported to display an unusual chromatin status in mouse trophoblast giant cells *in vivo* ^73^, albeit different from the one we described. This raises questions as to whether there are evolutionary advantages linked to XCI plasticity in extraembryonic tissues or whether this is merely reflecting the short-term life of these tissues, in which repressive locks such as DNA methylation are dispensable.

The most striking conclusion of our study is that one X chromosome must be silenced for female hESC-derived extraembryonic cell survival, which reveals a functional interplay between X chromosome dosage and extraembryonic cell fitness in humans. We provide direct evidence that *XIST* is required for *de novo* establishment of XCI in human models of early development. It is interesting to note that *XIST*-deficient hESCs can exit naïve pluripotency. This contrasts with mice, where this step is blocked by the presence of two active X chromosomes, which interfere with MAPK, Gsk3, and Akt signaling pathways ^74^. Such a difference is yet to be related to that of XCI kinetics: XCI is established concomitantly to pluripotency exit in mice ^75^, while in the system we used, it appears to be initiated between day 4 and 8 of differentiation, when cells have already reached an intermediate epiblast stage. Because *XIST*-deficient cells fail to achieve X-chromosome dampening ^21,26^, our results also suggest that dampening is not required for naïve pluripotency exit in humans, at least *in vitro*.

From scRNA-seq analyses, we concluded that differentiation could progress further toward extraembryonic fate even when XCI is blocked; by day 8 of differentiation, no differences in transcriptomic signatures, hence cell composition, could be observed between WT and *XIST*-deficient cells. The TSC and EXMC states can, however, not be maintained when both X chromosomes are active, and cells die within a few days. Several non-mutually exclusive hypotheses may explain the lag between XCI establishment (between day 4 and 8 of differentiation) and lethality of XCI-deficient cells (between day 10 and 15): there could be a delay between the increase in mRNA levels and its functional consequences on cellular homeostasis, which may involve the interaction of multiple pathways; appropriate dosage of X-linked products may be critical for differentiation events occurring later than day 8; and X-linked genes responsible for the phenotype may be inactivated later than the bulk of the chromosome. In any case, the phenotype we described is to be related to the requirement of X-chromosome dosage compensation for the development of extraembryonic ectoderm progenitors in mice ^13,14^, even if these cells have no equivalent in humans.

In conclusion, our work reveals the role of *XIST* for the establishment of XCI in humans and the requirement for XCI for female extraembryonic cell survival in humans, at least *in vitro*. It suggests that more subtle misregulation of XCI that may fortuitously occur during female human development could lead to embryo loss at peri-implantation stages due to failure in establishing placental progenitor programs, which remains to be investigated in embryos.

### Limitations of the study

We report the consequences of impaired XCI during *in vitro* differentiation using specific differentiation media; the effect of *XIST* loss in other differentiation conditions remains to be examined, also taking into consideration the fact that the efficiency of differentiation may vary between cell lines. The extent to which the phenotype would be reproduced in human embryo experiments remains to be determined, even if highly challenging. Indeed, naive hPSCs exhibit monoallelic *XIST* clouds, whereas the epiblast in pre-implantation embryos displays biallelic *XIST* clouds. The consequences of such a difference between the 2D stem cell model and the embryo warrant further investigation.

Additionally, a broader limitation within the naive hPSC field is the heterogeneity in chromatin landscapes arising from variations in culture conditions, which may impact the interpretation of findings.

In our *in vitro* system, TSCs and EXMCs exhibited varying degrees of XCI skewing. Skewing of XCI can arise through two mechanisms: primary and secondary. Primary skewing, where one X chromosome is preferentially inactivated during XCI initiation, and secondary skewing, which results from selection bias acting after random XCI has been established. However, it remains unclear whether the observed skewing of XCI in TSCs and EXMCs reflects primary or secondary skewing or a combination of both.

The *XIST* iKD cell lines enabled conditional *XIST* depletion, which was helpful in delineating the role of *XIST* during the differentiation; however, constitutive knockout of *XIST* could be employed in future studies.

Computational analyses of X chromosome expression upon *XIST* loss have some limitations: changes in the X/A ratio may result from differences in the process of X-chromosome upregulation; allelic analyses are limited to SNP-carrying genes, which are only a fraction of all X-linked genes.

## SUPPLEMENTAL FIGURE LEGENDS

**Figure S1:**
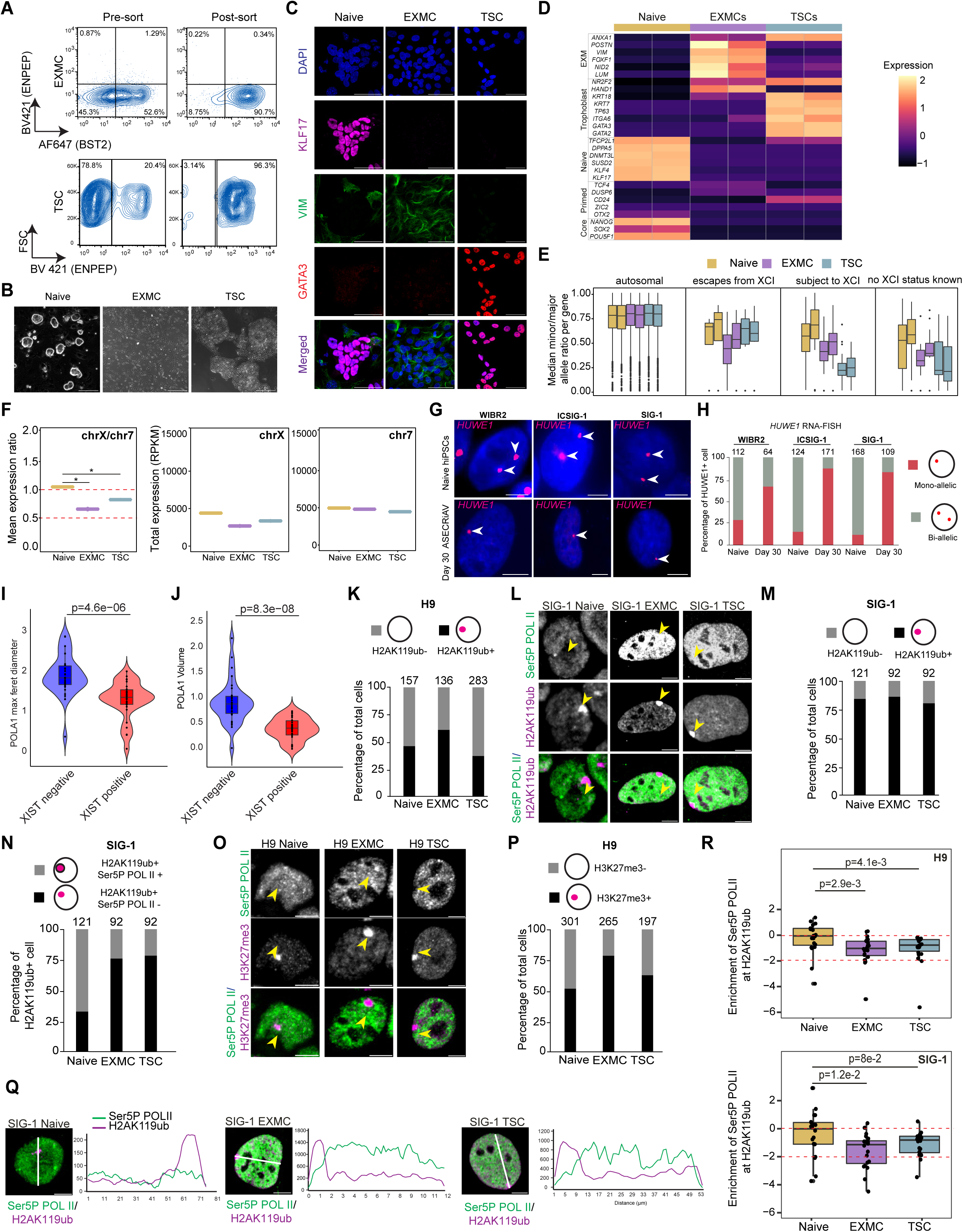
XCI is established during the differentiation of naive hESCs toward extraembryonic lineages, related to Figure 1. **A.** Flow cytometry contour plots showing both the BST2 positive and negative populations in both pre-sort and post-sort EXMCs (top) and the ENPEP positive and negative populations in both pre-sort and post-sort TSCs (bottom). **B.** Representative bright-field microscopy images showing the morphology of H9 naive hESCs, EXMC, and TSC. Scale bar, 1000 µm. **C.** Representative IF images showing lineage markers KLF17 (Magenta), GATA3 (Red), and VIM (Green) in H9 naive hESCs, EXMCs and TSCs. DAPI is shown in blue. Scale bar, 50 µm. **D.** Heatmap showing the expression of key marker genes for EXM, trophoblast, naive, primed, and core pluripotency genes in H9 naive hESCs, EXMCs and TSCs. Samples are plotted in the X-axis and genes are on the Y-axis. Expression is shown as a z-score calculated per gene. **E.** The median SNP ratio of the least expressed allele to most expressed allele of X chromosome in H9 naive hESCs, EXMC and TSCs split by previously published XCI status ^67^. There were 5851 informative genes on autosomes, 14 informative genes escaping XCI, 53 informative genes subject to XCI and 41 informative genes with no known XCI status. **F.** The mean X chromosome expression to mean chromosome 7 expression ratio in H9 naive hESCs, EXMCs and TSCs (left). Total X chromosome expression in RPKM in H9 naive hESCs, EXMCs and TSCs (Middle). Total chromosome 7 expression in RPKM in H9 naive hESCs, EXMC and TSCs (Right). **G.** Representative RNA FISH images of X-linked gene *HUWE1* (Magenta) in three different genetic backgrounds of naive and day 30 trophoblast medium. Scale bar, 50 µm. **H.** Bar plot showing the proportion of biallelic and monoallelic X-linked gene spots in three different genetic backgrounds of naive hPSCs and day 30 trophoblast cells with X-linked gene signals. The number of counted nuclei is indicated above each bar. **I.** Violin plots of maximum feret diameter (in µm) of POLA1 focal point in *XIST*-positive and *XIST*-negative alleles in day 0 H9 naive hESCs. Quantifications were performed on 31 nuclei from 2 batches of experiments. Each dot shows a unique nucleus. Statistical significance was assessed using a Wilcoxon rank-sum test for comparison between *XIST*-positive vs. *XIST*-negative alleles. P-values are shown in the figure. A p-value < 0.05 was considered statistically significant. **J.** Violin plots of maximum volume (in µm³) of POLA1 focal point in *XIST*-positive and *XIST*-negative alleles in day 0 H9 naive hESCs. Quantifications were performed on 31 nuclei from 2 batches of experiments. Each dot shows a unique nucleus. Statistical significance was assessed using a Wilcoxon rank-sum test for comparison between *XIST*-positive vs. *XIST*-negative alleles. P-values are shown in the figure. A p-value < 0.05 was considered statistically significant. **K.** Bar plot showing the quantification of the number of H2AK119ub foci in H9 naive hPSCs, EXMC, and TSC detected by IF. The number of counted nuclei is indicated above each bar. **L.** Representative IF images of Ser5P POL II (Green) and H2AK119ub (Magenta) in SIG-1 naive hiPSCs, EXMCs and TSCs. Scale bar, 5 µm. **M.** Bar plot showing the quantification of the number of H2AK119ub foci in SIG-1 naive hPSCs, EXMC, and TSC detected by IF. The number of counted nuclei is indicated above each bar. **N.** Bar plot showing the quantification of the number of H2AK119ub foci with or without Ser5P POL II in SIG-1 naive hiPSCs, EXMC, and TSC. The number of counted nuclei is indicated above each bar. **O.** Representative IF images of Ser5P POL II (Green) and H3K27me3 (Magenta) in H9 naive hESCs, EXMCs and TSCs. Scale bar, 5 µm. **P.** Bar plot showing the quantification of the number of H3K27me3 foci in H9 naive hESCs, EXMC, and TSC of the previous panel. The number of counted nuclei is indicated above each bar. **Q.** Left panel: Representative IF images of Ser5P POL II (Green) and H2AK119ub (Magenta) in SIG-1 naive hiPSCs, EXMCs and TSCs. Scale bar, 5 µm. Right panel: Representative line plots showing the gray intensity value of Ser5P POL II across a line passing through the H2AK119ub cloud. **R.** Box plot showing the quantification of the average gray intensity value of Ser5P Pol II at H2AK119ub foci as compared to outside of H2AK119ub foci in Naive, EXMCs and TSCs from panels 1L (for H9) S1L (for SIG-1). 20 random nuclei were chosen from each cell type for quantification. t-test p-values are given for the indicated comparisons. For panels H, K, M, N and P, the number of nuclei counted is indicated in supplemental table S1.

**Figure S2:**
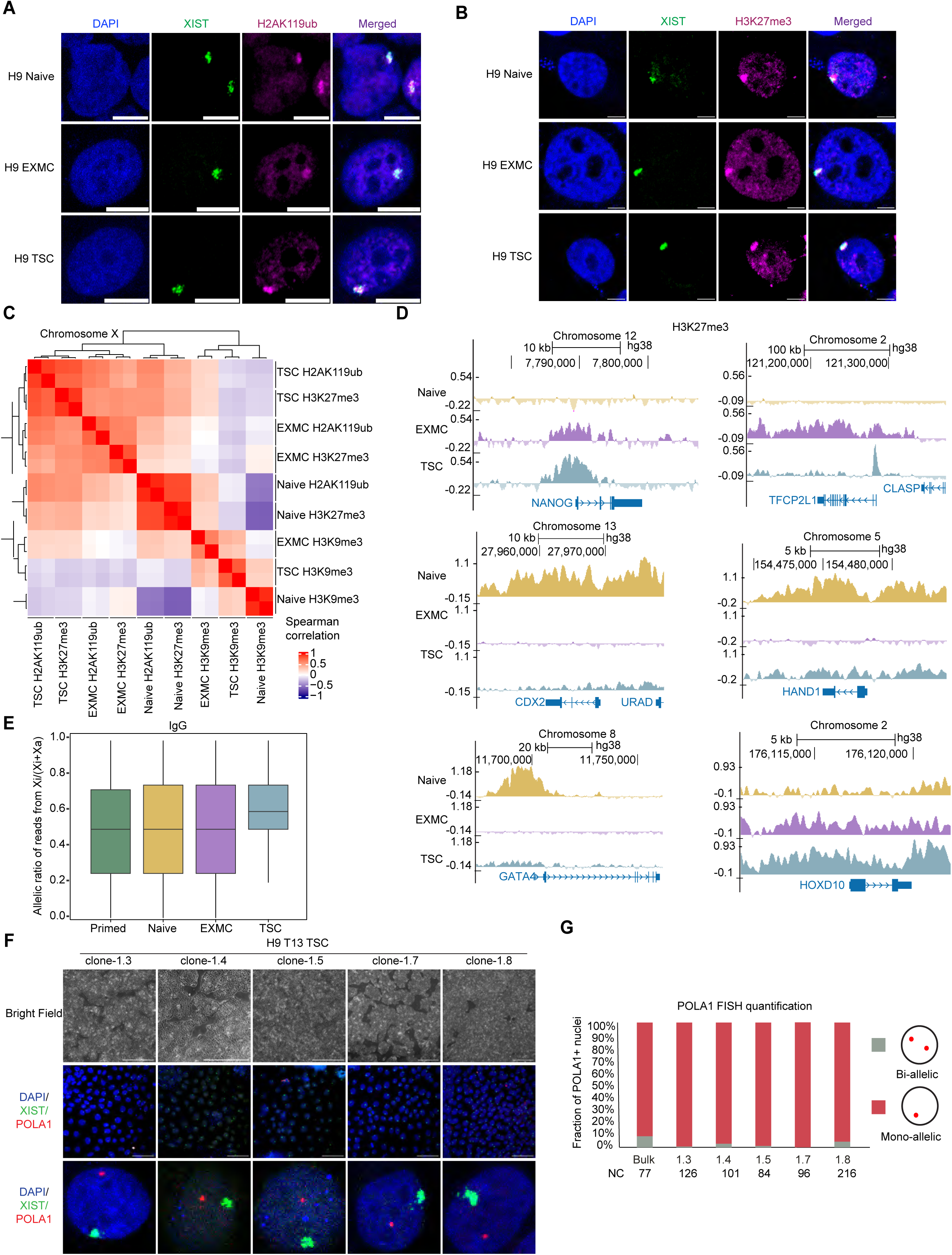
Unusual chromatin remodeling accompanies XCI during extraembryonic specification, related to Figure 2. **A.** Representative IF-FISH of *XIST* (Green) and H2AK119ub (Magenta) images showing co-localization of the repressive mark with *XIST* in H9 naive, EXMCs and TSCs. Scale bar, 5 µm. **B.** Representative IF-FISH of *XIST* (Green) and H3K27me3 (Magenta) images showing co-localization of the repressive mark with *XIST* in H9 naive, EXMCs and TSCs. Scale bar, 5 µm. **C.** Heatmap showing Spearman correlation coefficients of the CUT&RUN signals on the X chromosome between all samples in H9 naive hESCs, EXMCs and TSCs. **D.** H3K27me3 CUT&RUN tracks (log2 enrichment over IgG) around NANOG, TFCP2L1, CDX2, HAND1, GATA4, and HOXD10 in naive hESCs, EXMCs and TSCs. **E.** Box plot showing the allelic ratio of CUT&RUN reads from Xi/Xtotal for IgG in Naive, EXMCs, and TSCs. **F.** Representative brightfield images of various H9 TSC clonal lines (passage 13), each derived from a single cell (top panel). Scale bar: 1000 µm. Corresponding FISH images of the same clonal lines showing *XIST* (green) and *POLA1* (red) expression (middle panel). Scale bar: 50 µm. Cropped images of individual nuclei from the top panel, highlighting nuclear morphology (bottom panel). Scale bar: 5 µm. **G.** Bar plots showing the quantification of the proportion of cells with monoallelic (Red) and biallelic (Gray) POLA1 expression in the different clonal H9 TSCs of the previous panel. The number of nuclei counted is indicated in supplemental table S1.

**Figure S3:**
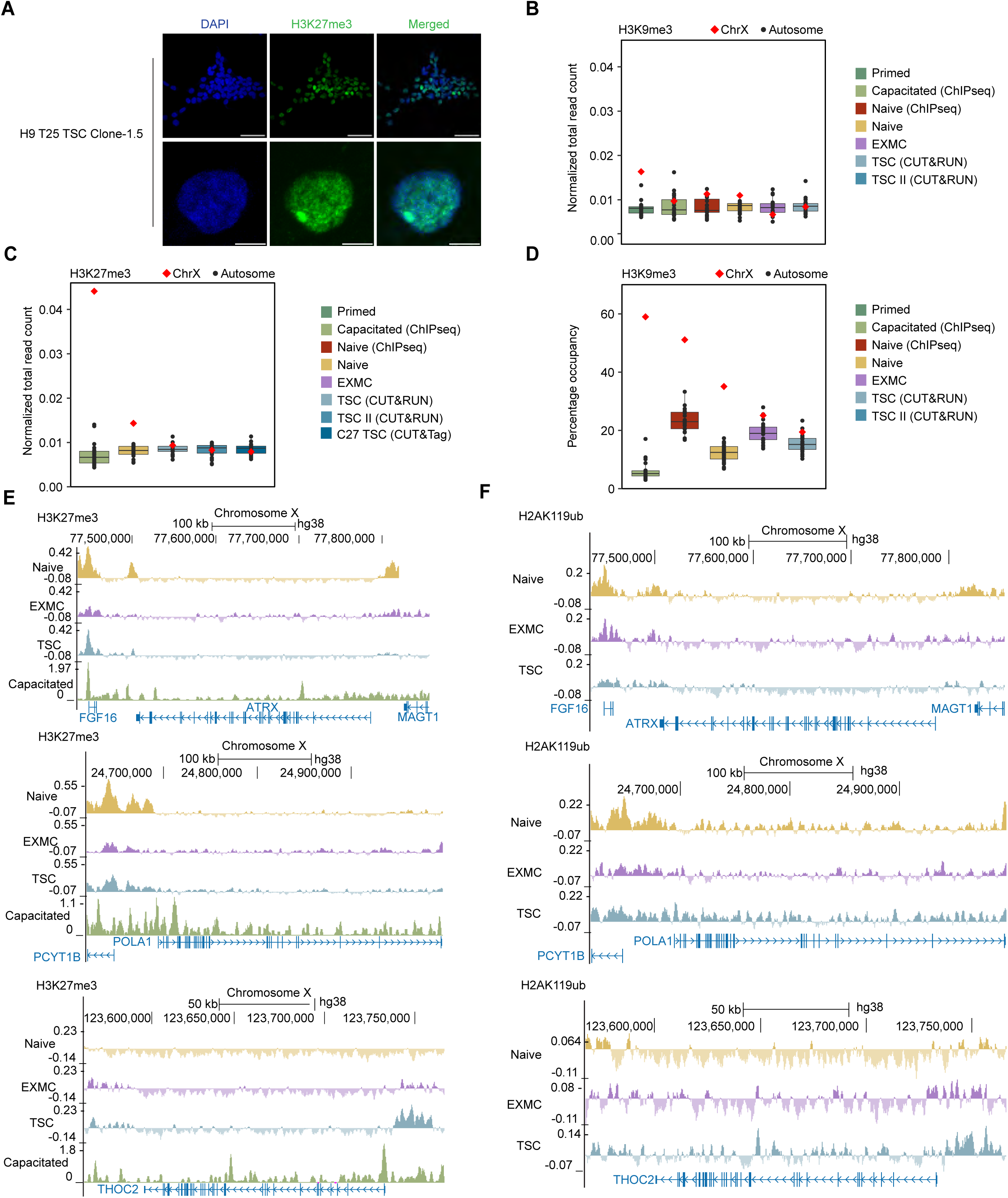
Unusual chromatin remodeling accompanies XCI during extraembryonic specification, related to Figure 2. **A.** Representative IF images of H3K27me3 (green) and DAPI (blue) in H9 TSC clone-1.5 (passage 25) (top panel). Scale bar, 50 µm. Cropped images of individual nuclei from the top panel, highlighting nuclear morphology (bottom panel). Scale bar: 5 µm. **B.** Total number of CUT&RUN H3K9me3 reads from each chromosome, normalized by the length of each chromosome in primed hESCs, capacitated hESCs (ChIPseq data), naive (ChIPseq data), naive hESCs, EXMCs and TSCs. **C.** Total number of H3K27me3 reads from each chromosome, normalized by the length of each chromosome in Capacitated cells hESCs (ChIPseq), naive hESCs, TSCs, H9 TSCs II (CUT&RUN) and CT27 TSCs (CUT&Tag). **D.** Percentage of H3K9me3 peaks occupancy per chromosome in capacitated cells hESCs (ChIPseq data), naive hESCs (ChIPseq data), H9 naive hESCs, EXMCs and TSCs. **E.** H3K27me3 CUT&RUN tracks (log2 enrichment over IgG) around the *ATRX*, *POLA1* and *THOC2* genes in naive hESCs, EXMCs and TSCs. **F.** H2AK119ub CUT&RUN tracks (log2 enrichment over IgG) around the *ATRX*, *POLA1* and *THOC2* genes in naive hESCs, EXMCs and TSCs.

**Figure S4:**
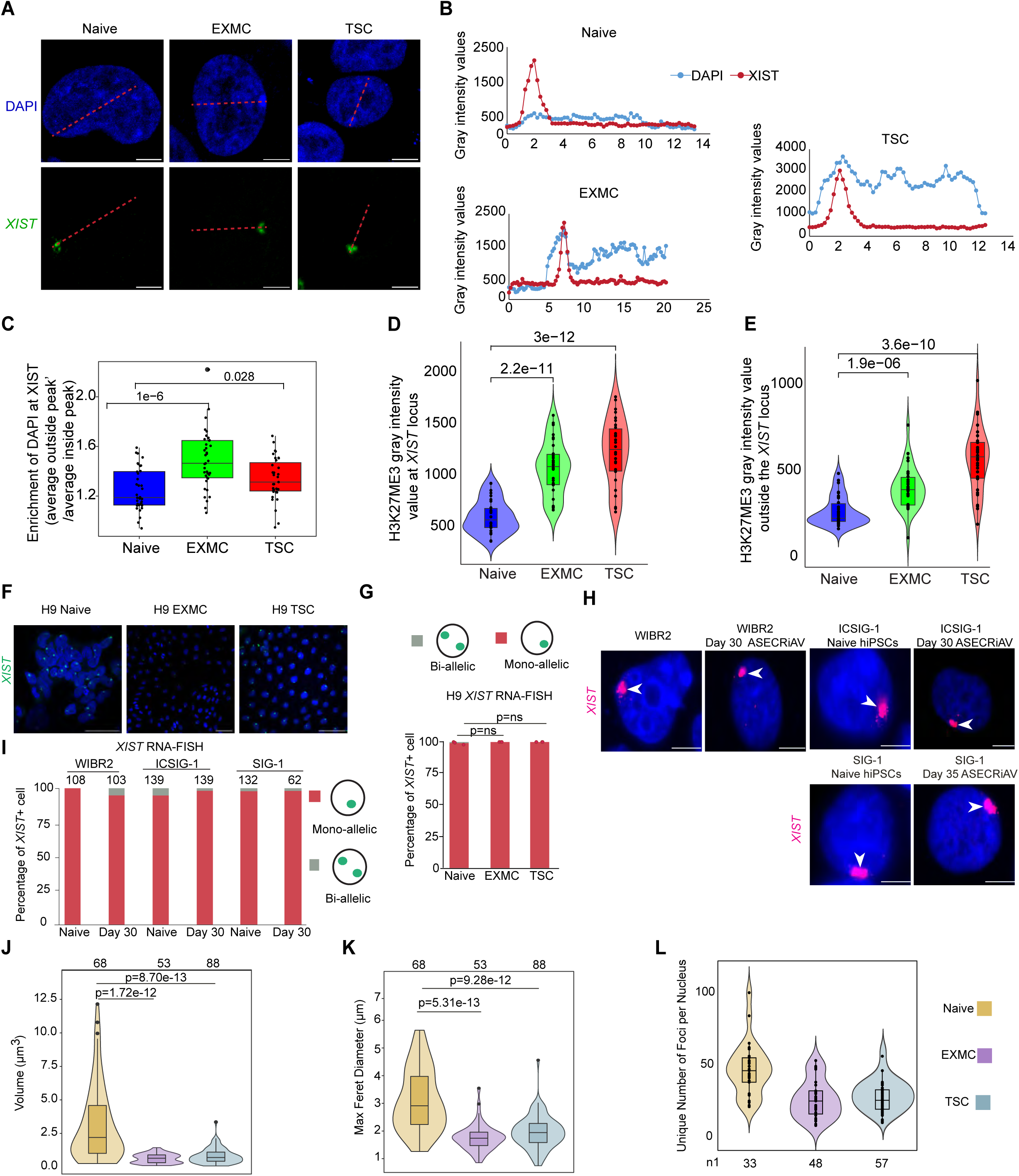
Unusual chromatin remodeling accompanies XCI during extraembryonic specification. related to Figure 2. **A.** Representative FISH images of DAPI (blue) and *XIST* (green) in H9 naive, EXMC and TSC. Scale bar, 5 µm. **B.** Representative line plots of one nuclei showing the gray intensity value of DAPI across a line passing through the *XIST* cloud. **C.** Box plot showing the quantification of the average gray intensity value of DAPI at *XIST* foci as compared to the outside of *XIST* foci in naive, EXMC, and TSC H9 from panel A. At least 30 random nuclei were chosen from each cell type for quantification. A student’s t-test was used to calculate significance. **D.** Violin plot showing the quantification of gray intensity value of H3K27me3 at *XIST* locus. At least 35 random nuclei were chosen from each cell type for quantification. Unless otherwise stated, statistical significance was assessed using a Wilcoxon test for comparisons between naive vs EXMC and naive vs TSC for each gene. P-values are shown in the figure. A p-value <0.05 was considered statistically significant. **E.** Violin plot showing the quantification of gray intensity value of H3K27me3 outside the *XIST* locus. At least 35 random nuclei were chosen from each cell type for quantification. Unless otherwise stated, statistical significance was assessed using a Wilcoxon test for comparisons between naive vs EXMC and naive vs TSC for each gene. P-values are shown in the figure. A p-value <0.05 was considered statistically significant. **F.** Representative RNA FISH images of *XIST* (green) in H9 naive hESCs, EXMCs and TSCs. Scale bar, 50 µm. **G.** Bar plots showing the quantification of the proportion of cells with either biallelic and monoallelic *XIST* clouds in H9 naive hESCs, EXMC, and TSC *XIST* RNA FISH, as shown in G. The number of nuclei counted is indicated below each bar for each round of experiments. Data represents the mean from 3 independent experiments (n = 3). Each experiment was performed in triplicate. Statistical significance was assessed using a logistic regression test. P-values are shown in the figure. A p-value <0.05 was considered statistically significant. **H.** Representative RNA FISH images of *XIST* (Magenta) in three different genetic backgrounds of naive hPSCs and day 30 trophoblast medium cells. DAPI is shown in blue. **I.** Quantification of the proportion of biallelic and monoallelic *XIST* clouds in three different genetic backgrounds of naive hPSCs and trophoblast medium cells with *XIST* FISH signals. The number of counted nuclei is indicated above each bar. **J.** Violin plots of maximum volume (in µm^3^) of *XIST* clouds in H9 naive hESCs, EXMCs and TSCs. The number of *XIST* positive nuclei counted is indicated above each condition. Wilcoxon-rank-sum p-values are given for the indicated comparisons. **K.** Violin plots of the maximum feret diameter (in µm) of *XIST* clouds in H9 naive hESCs, EXMC and TSCs. The number of *XIST* positive nuclei counted is indicated above each condition. Wilcoxon-rank-sum p-values are given for comparisons indicated by horizontal black lines. **L.** Representative violin plot showing the number of *XIST* foci per *XIST* cloud per nuclei in H9 naive hESCs, EXMCs and TSCs *XIST* RNA FISH, as shown in panel I. The number of nuclei counted is indicated below each violin plot.

**Figure S5:**
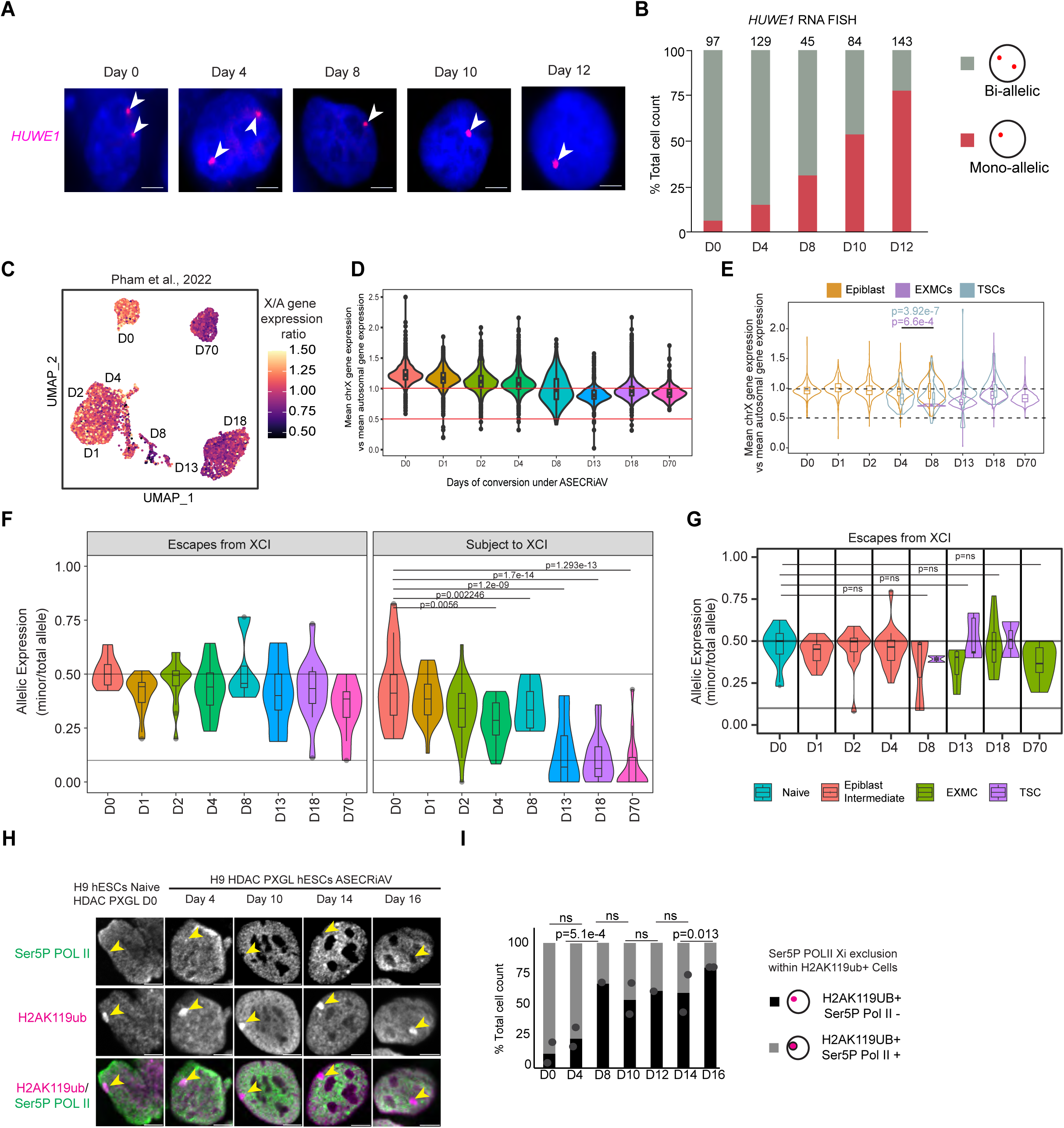
The establishment of XCI is closely linked with the acquisition of extraembryonic identity, related to Figure 3. **A.** Representative X-linked gene RNA FISH image showing *HUWE1* expression dynamics during H9 5iLA-PXGL naive hESCs to EXMCs and TSCs differentiation in ASECRiAV culture conditions. Samples were collected at five different time points as indicated. **B.** Bar plot showing the quantification of the proportion of *HUWE1* puncta during the differentiation of H9 5iLA-PXGL naive hESCs to EXMCs and TSC in the previous panel. The number of counted nuclei is indicated above each bar. **C.** Projection of the X to autosome gene expression ratio into the UMAP of time course scRNA-seq reanalysis of ^44^. **D.** Violin plots showing the X:A gene expression ratio of time course scRNA-seq reanalysis of ^44^. The data is shown with all cell types combined. **E.** Violin plots showing the X:A gene expression ratio of time course scRNA-seq reanalysis of ^44^ with epiblast, TSCs and EXMCs split apart. Statistical significance was assessed using a t-test. P-values are shown in the figure. A p-value <0.05 was considered statistically significant. **F.** Violin plot showing allelic expression of time-course scRNA-seq reanalysis of ^44^, calculated using the scLinaX pipeline ^65^. Data is presented with all cell types combined. Genes expected to escape XCI as well as genes expected to be inactivated are shown in Figure 3I. Statistical significance was assessed using a t-test. P-values are shown in the figure. A p-value <0.05 was considered statistically significant. **G.** Violin plot showing allelic expression of time-course scRNA-seq reanalysis of dataset ^44^, calculated using the scLinaX pipeline ^65^. Data is presented with data split by cell types. Only genes expected to be inactivated are shown. Statistical significance was assessed using a t-test. P-values are shown in the figure. A p-value <0.05 was considered statistically significant. **H.** Representative IF of Ser5P POL II (Green), H2AK119ub (Magenta) images showing dynamics during naive to TSCs-EXMCs conversion. Samples were collected at five different time points, as indicated. Scale bar, 5 µm. **I.** Representative bar plot showing the quantification of Ser5P POL II enriched nuclei during the differentiation of naive to TSC and EXMCs in the previous panel.

**Figure S6:**
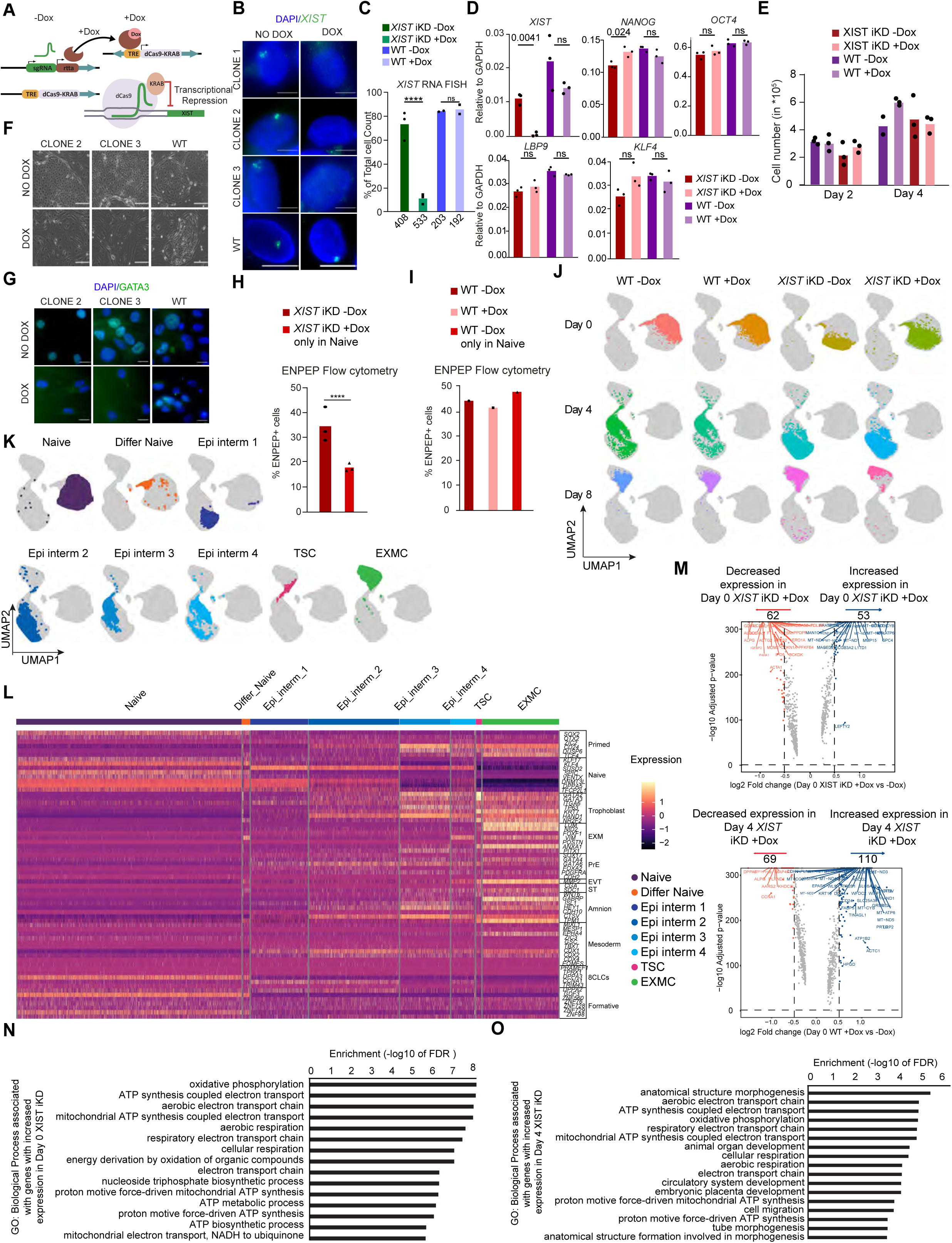
Loss of XIST dramatically impacts human extraembryonic cell survival, related to Figure 4. **A.** Schematic illustration showing the inducible CRISPR-Cas9 iKD system used to repress *XIST* expression in female H9 hESCs. **B.** Representative RNA FISH images of *XIST* (Green) in WT and *XIST* iKD clone-1, 2, 3 naive hESCs in PXGL culture media with or without Dox. Scale bar, 10 µm. Three independent numbers of *XIST* iKD clones (n=3) and two WT clones (n=2) were used for this experiment. **C.** Bar plot showing the quantification of the proportion of *XIST* clouds in cells of the previous panel; Chi² test was used when comparing -Dox vs +Dox conditions, resulting in P<0.0001 for the three independent iKD clones and P=0,1865 for the two WT clones (n=2). Each dot represents one of three independent, unique *XIST* iKD clones (n=3) and 2 WTclones (n=2). Statistical significance was assessed using a t-test. A p-value <0.05 was considered statistically significant. *** tests were significant beyond the limits (<e^-16^) of the test to calculate. **D.** Bar plot showing quantification of *XIST* RNA levels and of key pluripotency markers *NANOG, OCT4, KLF4,* and *LBP9* assessed by RT-qPCR. Statistical significance was assessed using unpaired t-test for comparisons between different conditions. P-values are shown in the figure. A p-value <0.05 was considered statistically significant. Three independent numbers of *XIST* iKD clones (n=3) and three WT clones (n=3) were used for this experiment. **E.** Cell proliferation was quantified for WT and *XIST* iKD naïve cells, comparing untreated versus 10 days of Dox treatment. Cells were seeded at the same density in each well and counted 2 or 4 days after passage. Each dot represents the average value of two measures taken per replicate clone. (n = 2-3 independent clones). **F.** Representative bright-field microscopy images showing the morphology of H9 *XIST* iKD clone-2, 3, and WT hESCs at Day 15 of trophoblast differentiation medium with or without Dox. Note: Feeders only are seen in the +Dox conditions. Scale bar, 200 µm. Three independent numbers of *XIST* iKD clones (n=3) and two WT clones (n=2) were used for this experiment. **G.** Representative IF of GATA3 (Green) of cells mentioned in S5E. DAPI is shown in blue. Scale bar, 20 µm. The experiment was performed in 3 independent *XIST* iKD clones of H9 hESCs, out of which two are shown here and one is shown in figure 4C (total n=3). **H.** Bar plots showing quantification of ENPEP positive cells by flow cytometry of Day 15 of ASECRiAV differentiation in *XIST* depleted and controlled cells. Chi² test was used to assess the statistical significance, resulting in P<0,0001. Each dot represents a unique *XIST* iKD clone (n=3 independent clones). **I.** Bar plots showing the quantification of ENPEP positive cells assessed by flow cytometry of Day 15 trophoblast medium culture in WT cells with Dox, without Dox and with Dox only in the naive state. Each dot represents a unique WT clone (n=1). **J.** UMAP plots from Figure 4E projecting individual conditions with their corresponding unique colors. **K.** UMAP plots from Figure 4F projecting individual cell types with their corresponding unique colors. **L.** Heatmap showing the expression of marker genes for primed pluripotency, core pluripotency, naive pluripotency, trophoblast and extraembryonic mesoderm (EXM), primitive endoderm (PrE), extravillous trophoblast (EVT), syncytiotrophoblast (ST), amnion, mesoderm, 8-cell-like-cells (8CLCs) and formative pluripotency. **M.** Differential gene expression as detected by scRNA-seq between *XIST* iKD cells with Dox vs without Dox, in Day 0 (top panel) and Day 4 (bottom panel). Dashed lines indicate -log10 adjusted p-Value <0.05 and log2 fold change < -0.5 or > 0.5. **N.** GO enrichment analysis of DEGs comparison between Day 0 *XIST* iKD cells with Dox vs without Dox is shown. **O.** GO enrichment analysis of DEGs comparison between Day 4 *XIST* iKD cells with Dox vs without Dox is shown.

**Figure S7:**
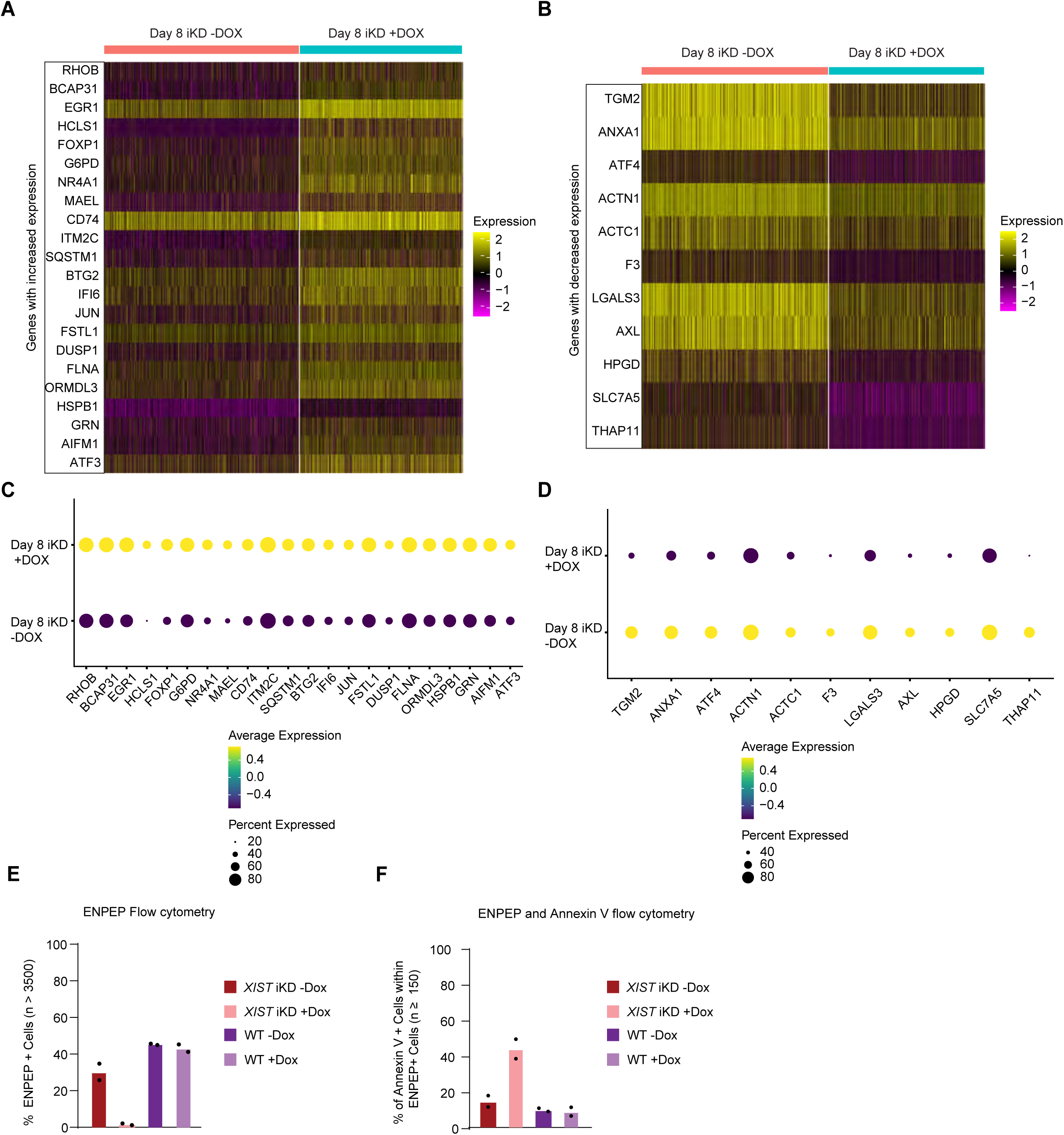
*XIST* is required for human extraembryonic cell fate survival, related to Figure 4. **A.** Heatmap showing increased expression of apoptotic DEGs associated with GO terms #1465, #1513, and #542 in day 8 *XIST* iKD cells, with or without *XIST*. **B.** Heatmap showing decreased expression of apoptotic DEGs with the same GO terms. **C.** Dot plot showing the expression patterns of all DEGs shown in panel A. **D.** Dot plot showing the expression patterns of DEGs from Panel B. **E.** Bar plot showing the relative proportion of ENPEP+ cells detected by FACS in WT and *XIST* iKD clones (+Dox, +Dox) at day 15 of ASECRiAV differentiation. 2 replicate clones were used per condition tested, collecting both attached and supernatant cells on the day of the experiment. **F.** Bar plot showing the percentage of Annexin V+ cells within the gated ENPEP+ cell population. 2 replicate clones were used per condition tested, collecting both attached and supernatant cells on the day of the experiment.

**Figure S8:**
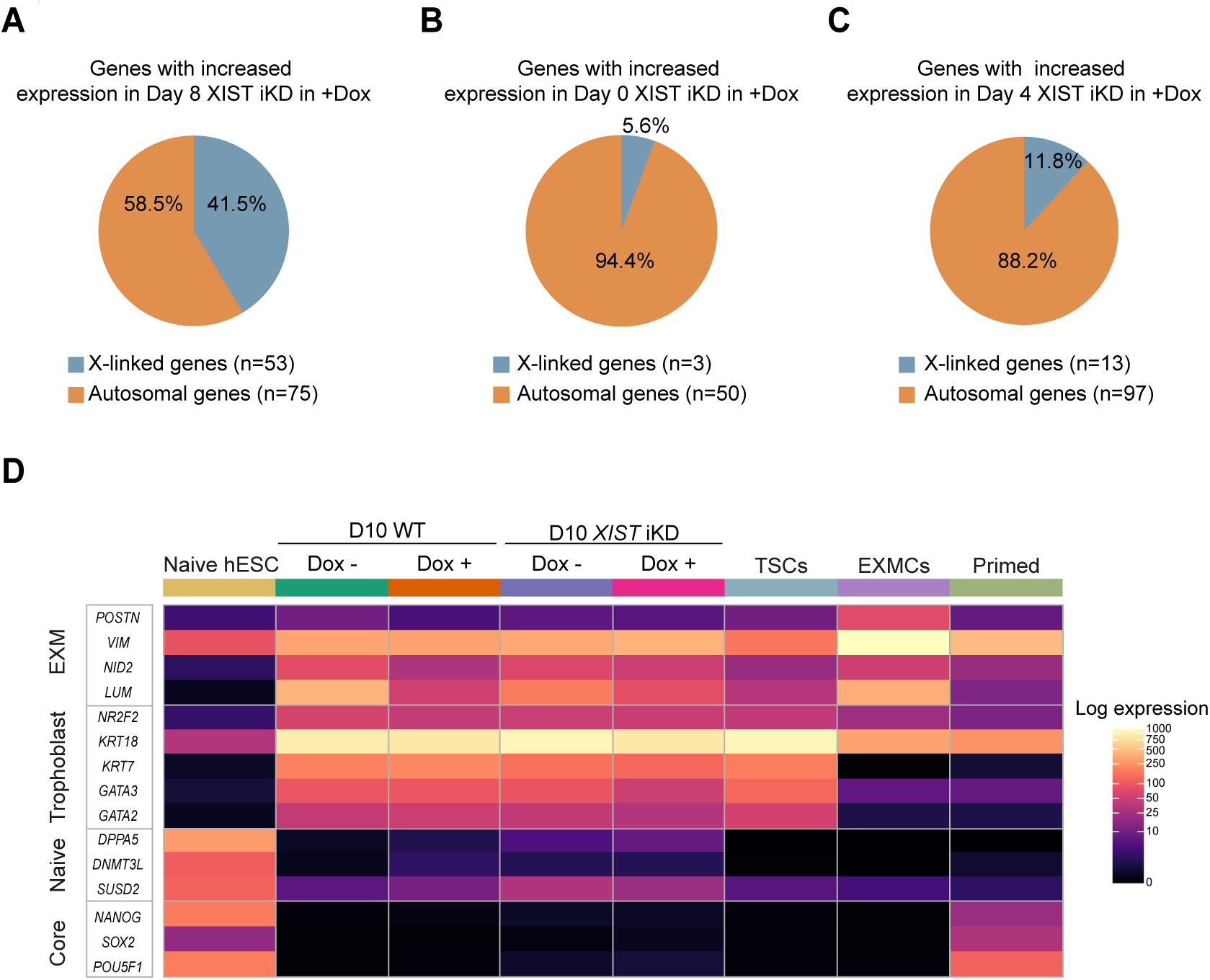
Human *XIST* is necessary for the establishment of human XCI in extraembryonic cells, related to Figure 5. **A.** Pie chart showing the proportion of DEGs in Day 8 *XIST* depleted cells compared to control (-Dox) cells corresponding to autosomal and X-linked genes. n is the number of genes. **B.** Pie chart showing the proportion of DEGs in Day 0 *XIST* depleted cells compared to control (-Dox) cells corresponding to autosomal and X-linked genes. n is the number of genes. **C.** Pie chart showing the proportion of DEGs in Day 4 *XIST* depleted cells compared to control (-Dox) cells belonging to autosomal and X-linked genes. n is the number of genes. **D.** Heatmap showing the log2 expression of key marker genes for EXM, trophoblast, naive, and core pluripotency in naive, day 10 WT -Dox, WT +Dox, *XIST* iKD -Dox and *XIST* iKD +Dox, TSCs, EXMCs and primed hESCs.

## Supplemental tables

**Supplemental Table S1: Number of nuclei counted for each round of FISH, IF, and IF-FISH experiments.**

**Supplemental Table S2: Number of cells at each annotation and time point in scRNA-seq time course, related to Figures 4, 5, S6, S7 and S8.**

**Supplemental Table S3: Differentially expressed genes found in the scRNA-seq time course, related to Figures 4, 5, S6, S7 and S8.**

**Supplemental Table S4: Gene ontology terms found related to differentially expressed genes in the scRNA-seq time course analysis, related to Figures 4, 5, S6, S7 and S8.**

**Supplemental Table S5: Differentially expressed genes separated by chromosome in the scRNA-seq time course analysis, related to Figures 4, 5, S6, S7, and S8.**

### Supplemental videos

**Video S1:** Reconstructed Z-stack 3D-SIM images of a segmented naive nuclei stained for XIST (Red), related to figure 1.

**Video S2:** Reconstructed Z-stack 3D-SIM images of a segmented EXMC nuclei stained for XIST (Red), related to figure 1.

**Video S3:** Reconstructed Z-stack 3D-SIM images of a segmented TSC nuclei stained for XIST (Red), related to figure 1.

## Key Resources Tables

### qPCR primers list

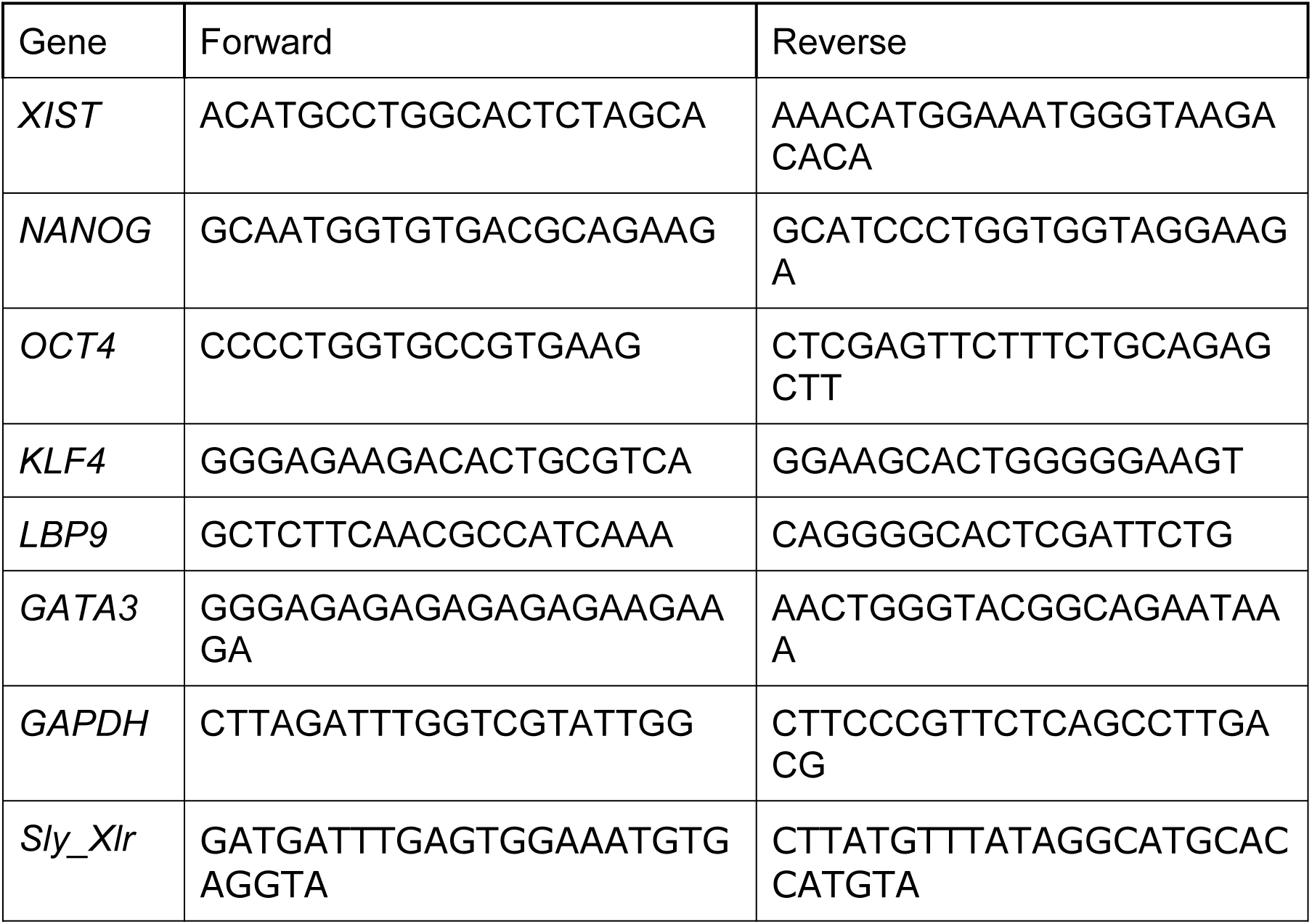

### CRISPRi sgRNA

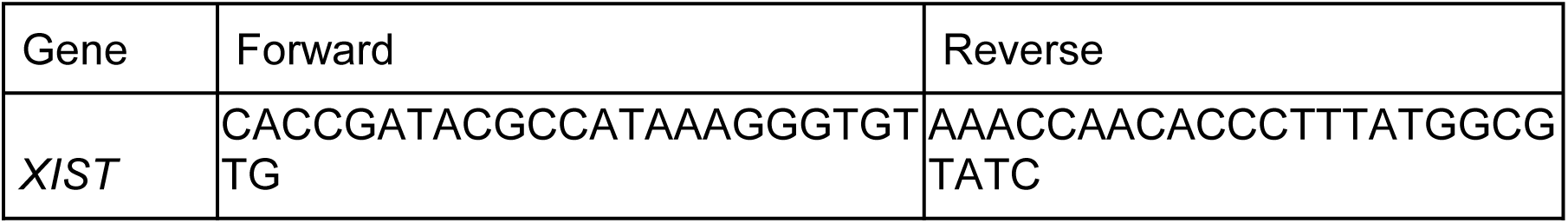

### IF antibodies list

| Reagent | Source | Identifier |
| --- | --- | --- |
| Rat monoclonal anti-GATA3 | Thermo Fisher | 14-9966-82 |
| Mouse monoclonal anti-Vimentin | Abcam | ab8978 |
| Rabbit polyclonal anti-KLF17 | Sigma-Aldrich | HPA024629 |
| Rabbit monoclonal anti-H2AK119ub | Cell Signalling | 8240s |
| Mouse monoclonal anti-H3K27me3 | Active motif | 61017 |
| Ser5P Polymerase II | Abcam | ab5408 |
| Annexin V | Invitrogen | 11-800572 |
| CD249 (ENPEP) | BD Biosciences | 744872 |
| Annexin V Apoptosis Detection Kit FITC | Invitrogen | 88-8005-72 |

### CUT&RUN antibodies list

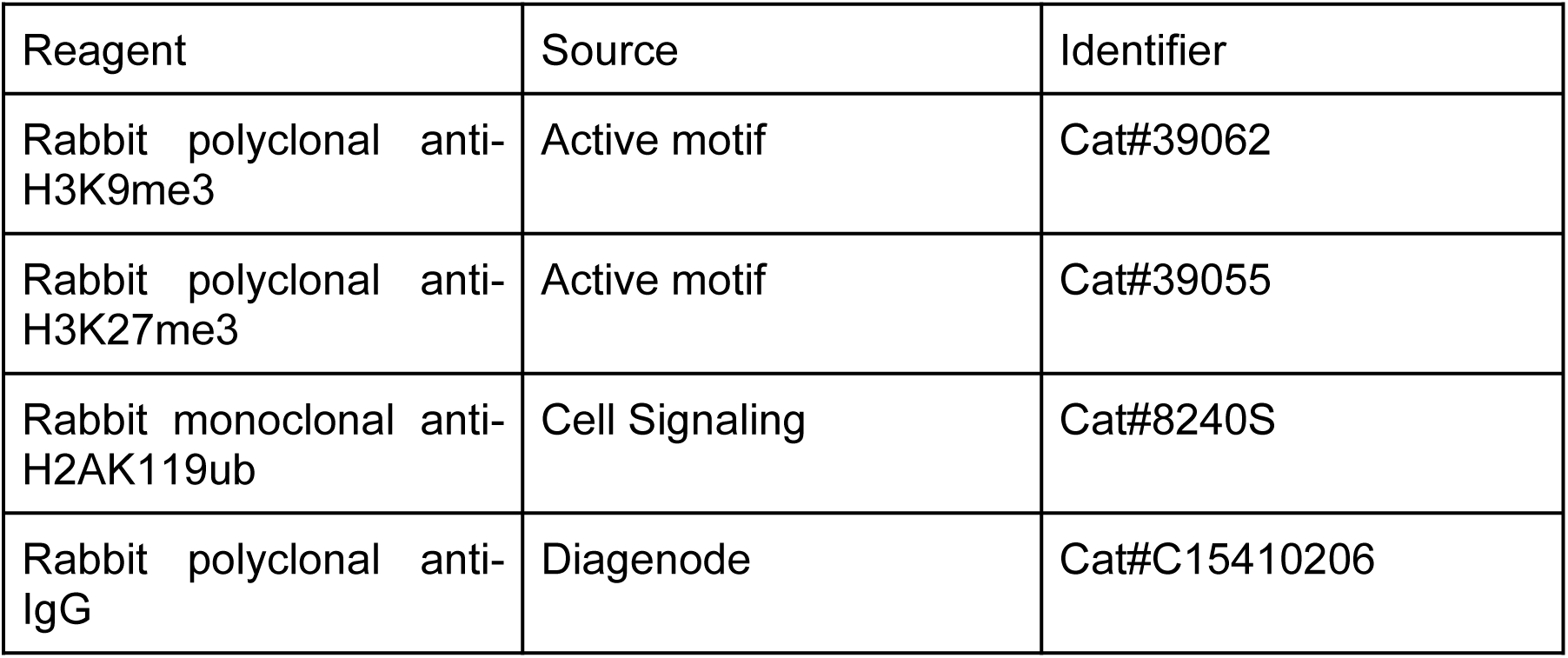

| Reagent | Source | Identifier |
| --- | --- | --- |
| Rabbit polyclonal anti-H3K9me3 | Active motif | Cat#39062 |
| Rabbit polyclonal anti-H3K27me3 | Active motif | Cat#39055 |
| Rabbit monoclonal anti-H2AK119ub | Cell Signaling | Cat#8240S |
| Rabbit polyclonal anti-IgG | Diagenode | Cat#C15410206 |

### Resource availability

#### Lead contact

Further information and requests for resources and reagents should be directed to and will be fulfilled by the lead contact, Claire Rougeulle and Vincent Pasque.

#### Materials availability

All stable reagents generated in this study are available from the lead contact without restriction except for human embryo derived cell lines and human induced pluripotent stem cell lines and their derivative for which permission must be requested from WiCell, Sigma or Dr. Rudolf Jaenisch and a material transfer agreement and other documents required by local legislation and institutional policies must be completed.

#### Data and code availability

Raw and processed sequencing data (scRNA-seq, bulk RNA-seq, CUT&RUN and MeD-seq) have been submitted to the NCBI GEO under accession number GSE261711.

This paper analyzes eXISTing, publicly available data. The accession numbers for the datasets are listed in the key resources table. All analysis code is available at: https://github.com/pasquelab/Human-XIST-XCI_AP.

Any additional information required to reanalyze the data reported in this paper is available from the lead contact upon request.

### Experimental model and subject details

#### Ethics

Work with human embryonic and induced pluripotent stem cells to model early human development was approved by the UZ/KU Leuven ethics committee (S52426, S64962, S66185, S66184 and S66375) and also by the Flemish Government (SBB 219 2020/0435). Work with mouse embryo for mouse embryonic fibroblast was approved by the UZ/KU Leuven ethics committee (P170/2019).

Experiments were carried out using the female H9 hESCs obtained from the WiCell Research Institute. Research on human embryonic stem cells has been approved by Agence de la Biomédecine and informed consent was obtained from all subjects.

### Cell lines

#### Human primed pluripotent stem cell culture

Human primed pluripotent stem cells were cultured as previously described in ^44^. H9 hESCs (WiCell#WA09), Sigma hiPSCs (Sigma#iPSC EPITHELIAL-1-IPSC0028, SIG-1) inducible Cas9 Sigma hiPSCs (Sigma#iPSC EPITHELIAL-1-IPSC0028, ICSIG-1) and WIBR2 29M-GP26-TN9 hESCs ^76^ were grown with or without feeder conditions, in normoxia conditions (5% CO2), and under humidified conditions at 37°C. In feeder-free conditions, cells were cultured in pre-coated geltrex (Thermo #A1413302) tissue culture treated plates in complete E8Flex medium (Stemcell technologies). Cells were dissociated into smaller clumps every 5-6 days by incubating 5 minutes at room temperature in Versene. In feeder-dependent conditions, primed hESCs were grown on 0.1% gelatin coated plates with mitomycin-treated mouse embryonic fibroblast (MMC-MEF) feeders in human knockout serum replacement (KSR) primed medium containing 77.5% of DMEM/F12 (Gibco, 31330-038), 15% FBS (Gibco, 10270106), 5% KSR (Gibco, 10828028), non-essential amino acid (Gibco, 11140050), 2 mM L-glutamine (Gibco, 25030081), Penicillin-streptomycin (Gibco, 15140-122), 0.1 mM β-mercapto-EtOH (Gibco, 31350-010) and adding 10ng/ml FGF2 (Peprotech) freshly every day. Cells were passaged every 6-7 days using a 20 minutes incubation in collagenase (ThermoFisher 17104019). Media was changed every day.

##### *XIST* iKD clonal lines

A sgRNA targeting the *XIST* promoter was designed using the web-based tool CRISPOR (http://crispor.tefor.net/) and cloned into the PB_rtTA_BsmBI vector (Addgene, #126028) ^77^. The sgRNA sequence can be found in the resource table. Using the Amaxa 4D-NucleofectorTM system (Lonza), one million H9 primed hESCs were transfected with 1.5 μg of PB_tre_dCas9_KRAB (Addgene, #126030), 0.75 μg of PB_rtTA_BsmBI and 1.5 μg of piggyBac transposase ^78^. Cells were then treated with G418 (350 μg/mL) and hygromycin (350 μg/mL) until separate colonies were obtained. The number of random insertions in the genome was verified by qPCR and clones with the lowest insertion number (n=2) were used for further experiments. Three independent primed H9 hESCs clones (named clone 1, clone 2, clone 3) and one WT clone (that has not been transfected) were then used.

#### Human primed to naive resettings

##### PXGL HDAC resetting

WT and *XIST* iKD naive H9 hESCs were generated by chemical resetting of the H9 primed hESCs using a protocol previously described with small modifications ^39,79^, and cultured on MMC-MEF. PXGL naive hESCs were cultured in PXGL medium consisting of a 1:1 mixture of DMEM/F12 (Thermo #31330038) and Neurobasal media (Thermo #21103-049) supplemented with 0.5% N2 supplement (#17502048), 1% B27 supplement (#17504044), 2mM l-glutamine (#25030081), 100 μM β-mercaptoethanol (#31350-010)l and 1x penicillin–streptomycin (#15140-122) (all from Gibco, Thermo Fisher Scientific) as well as 1μM PD0325901 (Axon Medchem, 1408; CAS: 391210-10-9), 2μM XAV939 (Cell guidance systems, SM38-10; CAS: 284028-89-3), 2μM Gö6983 (Tocris, 2285; CAS: 133053-19-7), 10 ng/mL human LIF (Peprotech, 300-05). Naive hESCs were routinely splitted as single cells every three days at a ratio of 1:3 using Accutase (Stemcell Technologies, 07920) and plated in fresh media supplemented with 10 μM of Y-27632 (Stemcell Technologies). For *XIST* depletion, *XIST* iKD hESCs were treated with 1μg/mL Dox for 10 days then either treated or untreated for subsequent culture with the same concentration of Dox.

##### KLF4 mRNA resetting

The resetting of human primed hESCs or hiPSCs was performed as previously described in ^44^. In summary, starting from day 1 or day 2 after seeding primed hESCs in E8 onto Geltrex, cells were lipofected daily with KLF4 mRNA (Miltenyi, 130-101-115) for 9 days ^80^. 2 μl KLF4 mRNA were diluted in 250 μL Opti-MEM (Gibco, 31985-047) and 6 μl of lipofectamine RNAimax (Invitrogen, 13778075) in another 250 μL of Opti-MEM for a single well of a 6-well plate. Then, the diluted mRNA was added to the diluted lipofectamine and incubated for 20 minutes at room temperature. After the medium was exchanged for 1.5 mL of fresh E8 flex, the mixture was added dropwise. After 10 days of lipofection, cells were passaged with Versene onto MMC-MEF feeders in E8 flex and transfected again. Starting from the following day, the medium was switched to t2iLGö (see below) supplemented with 10 mM Y-27632 (Tocris, 1254), and cells were transferred to hypoxia, while the lipofection was repeated every day for another 5 days. The naive t2iLGö medium contains a 50:50 mixture of DMEM/F12 (Gibco, 31330-038) and Neurobasal medium (Gibco, 21103-049), supplemented with 2 mM L-glutamine (Gibco), 0.1 mM β-mercaptoethanol (Gibco, 31350-010), 0.5% N2 supplement (Gibco, 17502-048), 1% B27 supplement (Gibco, 17504-044), 1% Penicillin-streptomycin (Gibco, 15140-122), 10 ng/ml human LIF (PeproTech, 300-05), 250 *μ*M L-ascorbic acid (Sigma-Aldrich, A4544-100G), 10 *μ*g/ml recombinant human insulin (Sigma, I9278-5ML), 1 *μ*M PD0325901 (Axon Medchem, 1408), 1 *μ*M CHIR99021 (Axon Medchem, 1386), 2.5 *μ*M Gö6983 (Tocris, 2285) ^80^. After a total of 15 days of transfection, cells were passaged with Accutase (Sigma-Aldrich, A6964-100ML) on the following day onto fresh feeders and continued to be cultivated in PXGL.

### 5iLA resetting

To convert primed hESCs to naive hESCs, trypsinized primed hESCs were seeded onto gelatin-coated MMC-MEF feeders tissue culture treated 6-well plates and cultured with human KSR primed medium along with 10 μM of Y-27632 (Tocris, 1254) in a humidified normoxia incubator (5% CO2) for two days. On the third day and after giving a wash with Phosphate-buffered Saline (PBS (Gibco, 10010-015)) the medium was changed to 5iLA medium composed of 1:1 DMEM/F12 (Gibco, 31330-038) and Neurobasal (Gibco, 21103-049), 1% N2-supplement (Gibco, 17502-048), 2% B27 supplement (Gibco, 17504-044), 20 ng/ml recombinant human LIF (PeproTech, 300-05), 2mM L-glutamine (Gibco, 25030-081), 1% non-essential amino acid, 0.1 mM β-mercaptoethanol (Gibco, 31350-010), 1x Penicillin-streptomycin (Gibco, 15140-122), 50 μg/ml BSA (Sigma-Aldrich, A3059) and supplemented with 5 inhibitors: PD0325901 (Stemgent, 1 μM), IM-12 (Enzo, 1 μM), SB590885 (R&D systems, 0.5 μM), WH-4-023 (A Chemtek, 1 μM), Y-27632 (Tocris, 10 μM), and Activin A (Peprotech, 20 ng/ml) and grown in a humidified incubator in hypoxia condition (5% CO2 and 5% O2) at 37°C. After an initial wave of cell death around day 10-13, dome-shaped naive colonies started appearing. These cells were passaged into single cells every 4-5 days using 5 minutes incubation in Accutase (Sigma-Aldrich, A6964-100ML) at 37°C.

For some experiments, H9 and WIBR2-MGT naive hESCs that were derived and cultured in 5iLA conditions were switched at passage 12 into PXGL naive medium for stable maintenance and expansion.

#### Human naive pluripotent stem cell culture

Naive hESCs (H9 hESCs, WIBR2-MGT (converted from WIBR2 29M-GP26-TN9 hESCs) and Sigma hiPSCs) were cultured on MMC-MEF feeders in 5% O2 and 5% CO2 incubators under humidified conditions at 37°C as previously described in ^44^. All naive hESCs were cultured in the PXGL medium. Naive hESCs were passaged every 4-5 days at a ratio of 1:2 or 1:3 by single-cell dissociation with Accutase (Sigma-Aldrich, A6964-100ML), followed by filtering through a 40 μm cell strainer (Corning, 352340).

#### Mouse feeders

MEFs were isolated from E14.5 pregnant WT C57/Black 6 mice. Male embryos were selected based on sex genotyping PCR and immortalized with Mitomycin C (Bioconnect). MEFs were cultured and harvested in a humidified incubator at 37°C and in 5% CO2 by using filter-sterilized MEF medium consisting of 90% DMEM (Thermo #41966-029) supplemented with 10% FBS (Thermo #10270106), 1% Glutamax (Thermo #35050061), 1x Penicillin-streptomycin (Thermo #15140-122), 1x non-essential amino acid (Thermo #11140050), and 0.1 mM β-mercaptoethanol) and on 0.1% gelatin-coated tissue culture-treated plates.

#### Naive human pluripotent stem cells to EXMC and TSC fate induction

Human naive to EXMC and TSC differentiation were carried out using the following previously described protocols for TSCs ^42,44–48^. Naive hESCs were seeded in such a way that expecting at least 70-80% confluency by Day 2 on MMC-MEFs after dissociating in single cells using TrypLE (Thermo Fisher, 12605-010) for 15 minutes at 37°C in their respective naive culture conditions supplemented with 10μM Y-27632 (Tocris, 1254). The next day, after a wash with 1X PBS (Gibco, 10010-015), the media was switched from naive to ASECRiAV medium ^42^, consisting of DMEM/F12 (Gibco, 31330038) supplemented with 0.3% BSA (Sigma, A3059), 0.2% FBS (Gibco, 10270-106), 1% Penicillin-streptomycin (Gibco, 15140-122), 1% insulin-transferrin-selenium-ethanolamine-X 100 supplement (Gibco, 51500056), 1.5 μg/ml L-ascorbic acid (Sigma, A8960), 0.5 μM A83-01 (Peprotech, 9094360), 1μM SB431542 (Axon Medchem, 1661), 50ng/ml hEGF (Miltenyi Biotec, 130-097-750), 2 μM CHIR99021 (Axon Medchem, 1386), 0.8 mM Valproic acid (Sigma, V0033000), 0.1 mM β-mercapto-EtOH (Gibco, 31350-010) and 5 μM Y-27632 (Tocris, 1254) or ACE medium^47^ consisting of 1:1 DMEM/F12 and Neurobasal, 0.5% N2-supplement, 1% B27-Supplement, 2 mM L-Glutamine, 0.1 mM β-mercaptoethanol, 1x penicillin-streptomycin, 1 μM of A83-01 (Peprotech, 9094360), 2 μM CHIR99021 (Axon Medchem, 1386), 50ng/ml hEGF (Miltenyi Biotec, 130-097-750) and 5 μM Y-27632 (Tocris, 1254). The medium was changed every two days supplemented with 5μM Y-27632. From passage one onwards, both EXMCs and TSCs were cultured and maintained on 5 μg/ml Collagen IV or iMatrix 511 silk (Laminin-E8) coated tissue culture-treated plates in hypoxia conditions (5% O2 and 5% CO2) and passaged every 5 days at 1:3 or 1:6 splitting ratios. Collagen IV-coated cell culture plates were coated overnight at 37°C and iMatrix 511 silk coated plates were coated for 1h.

#### *XIST* iKD naive human pluripotent stem cells to EXMC and TSC fate induction

Naive *XIST* CRISPRi hESCs were cultured in PXGL medium supplemented with 1 µg/ml doxycycline for 10 days to ensure *XIST* transcriptional repression. The differentiation into EXMCs and TSCs was carried out as previously described, with minor modifications: naive hESCs were dissociated in single cells using Accutase (Stemcell, 07920) for 5 minutes at 37°C. 15.625 cells/cm^2^ were plated on MMC-MEFs in PXGL supplemented with 10μM Y-27632 (Tocris, 1254). The next day, after a wash with 1X PBS (Gibco, 10010-015), the media was switched from PXGL/PXGL+Dox to ASECRiAV/ASECRiAV+Dox medium ^42^. The medium was changed every day and cells were passaged every 5 days at 1:3 dilution, plated on MMC-MEFs.

### Method details

#### Immunofluorescence

Immunofluorescence staining was performed as described previously ^44,81^ with little modification. Cells were grown on 0.1% gelatinized 18-mm round coverslips with or without feeders. The next day, cells were fixed in 4% paraformaldehyde (PFA) diluted in 1X PBS for 10 minutes at room temperature in the dark, permeabilized with 0.5% Triton X-100 in PBS for 10 minutes and washed twice with 0.2% Tween20 in PBS (PBST) for 5 minutes each before proceeding to the staining. After this step, cells were either stored at 4°C or directly subjected to staining. However, we noted that the H3K27me3 does not last long at 4C after fixation. Primary and secondary antibodies were diluted in a blocking buffer containing mainly PBST with 5% normal donkey serum and 0.2% fish skin gelatin. A list of antibodies can be found in the resource table. Cells on coverslips were incubated at 4°C with the specific primary antibodies in blocking solutions (1:100 dilution for all antibodies except 1:50 dilution for NANOG, 1:40 dilution for FOXA2), and after that washed three times with PBST each 5 min. After that samples were incubated with the appropriate corresponding fluorophore conjugated secondary antibodies in blocking buffer (1:500 dilution) for 1h in the dark, washed 2 times with PBST 5 minutes each, washed with 0.002% DAPI (Sigma-Aldrich, D9542) solution in PBST and a last wash again with PBST for 5 minutes. The coverslips were mounted in Prolong Gold reagent with DAPI after a final wash in PBST. Mounted coverslips were kept at room temperature in the dark overnight before imaging.

#### RNA FISH and RNA FISH combined with IF

RNA FISH was performed as described previously ^81^ with little modification. Cells were grown on 18-mm round coverslips with their respective adhesive matrix or feeders. The next day, cells were transferred to fresh 12-well plates after a series of washes with 1X PBS containing CaCl2 and MgCl2. Cells were then fixed in 4% paraformaldehyde-PBS for 10 minutes at room temperature in the dark. After washing the coverslips with PBS to remove the remaining PFA, cells were permeabilized with cold 0.5% Triton X-100 in PBS for 10 minutes and washed twice with cold 0.2% Tween20 in PBS (PBST) for 5 minutes each. After this step, cells were either stored at -20°C in 70% ethanol or directly subjected to RNA FISH or IF-FISH staining. When combining IF with RNA FISH, IF was performed as described above with the addition of RNAaseOUT (Invitrogen) and tRNA (Invitrogen) to the blocking buffer in all steps. After IF, cells were re-fixed in 4% paraformaldehyde, followed by 2X SSC. Cells were kept at least for 1-2 hours at 4°C in 70% ethanol before starting FISH. RNA FISH for *XIST* was performed with DNA-modified RNA probes labeled with ChromaTide Alexa Fluro 488-5-DUTP (Thermo C11397), ChromaTide Alexa Fluro 594-5-DUTP (Thermo C11400) or Green 496 dUTP (Enzo ENZ-42831L) and AMINOALLYL DUTP CY(R)5 (Sigma 19475) using the Bioprime kit (Thermo 18095012) from a plasmid containing *XIST exon-1* ^82^, and X-linked genes were obtained from BAC containing *HUWE1* (Bacpac resource center, RP11-975N19), *POLA1* (Bacpac resource center, RP11-11104L9), *ATRX* (Bacpac resource center, RP11-42M11), and *THOC2* (Bacpac resource center, RP11-121P4). When performing any FISH experiments, all the solutions contained 2mM of the ribonucleoside-vanadyl complex. Before the hybridisation steps, coverslips were serially diluted with 70%-100% ethanol. RNA probes were pre-hybridized at 37°C at least for 1h before use. RNA probes and samples were hybridized overnight at 37°C. Afterwards samples were washed with 3 rounds of SSC solutions (1:1 formamide-2X SSC, 2X SSC and 1X SSC) at 42°C by incubating 5-15 minutes each. Finally, the coverslips were mounted on the glass slides using Prolong Gold Antifade Mountant (Thermo P36930). Mounted coverslips were kept at room temperature in the dark overnight before imaging.

#### Microscopy

Imaging of WT naive hESCs, EXMCs and TSCs was done with a Zeiss Axioimager A1 inverted microscope with an AxioCam MRc5 camera, Nikon TiA1R confocal microscope and processed in ImageJ. FISH and IF images were taken using a Nikon TiA1R confocal microscope. Bright field images were taken using a Nikon Eclipse Ti2 microscope and analyzed using ImageJ software.

Imaging of XIST iKD clone experiments was done on DMI-6000 inverted microscope with a motorized stage (Leica), equipped with a CCD Camera HQ2 (Roper Scientifics) and an HCX PL APO 64-100X oil objective (numerical aperture, 1.4, Leica) using the Metamorph software (version 7.04, Roper Scientifics). Brightfield images were taken using a Leica DM IL LED microscope equipped with a C4742-98-24ERG digital camera (Hamamatsu).

##### RNA FISH imaging

40 optical z-sections were acquired at 0.3µm at different wavelengths depending on the modified-dUTPs: DAPI (360 nm, 470 nm), FITC (470 nm, 525 nm), Cy3 (550 nm, 570 nm), and Cy5 (647 nm, 668 nm). Stacks were processed using ImageJ and are represented as a 2D stacked maximum projection.

##### Super-resolution microscopy

3D-structured illumination microscopy (3D-SIM) was performed on an inverted Zeiss Elyra 7 (with lattice SIM module for structured illumination) microscope equipped with two PCO.edge 4.2 CLHS sCMOS cameras in combination with a 63X oil objective (NA 1.4). Alexa Fluor 555 was excited with a 561 nm green diode with 500 mW laser power and emission was collected with a filter allowing 570-620 nm emission detection. DAPI was excited with a 405 nm diode with 50 mW laser power and emission was collected with a filter allowing 420-480 nm emission detection. Raw images were computationally reconstructed using ZEN black (software version, Carl Zeiss Microscopy GmbH) and subsequently imported into ImageJ/Fiji or IMARIS software.

##### Super-resolution microscopy image analysis

3D-SIM image stacks were converted from .czi to .ims format using Imaris Converter 10.2.0 (Oxford Instruments) and analyzed in Imaris 10.2.0 using the Cells tool for segmentation. Nuclei were segmented based on the DAPI channel, while *XIST* foci were detected from the *XIST* channel using the vesicle detection pipeline within the Cells tool.

Nuclear segmentation was performed using an intensity threshold of 20 and a minimum object diameter of 10 μm. *XIST* foci were identified using the spot detection feature, with a minimum estimated diameter of 0.15 μm and a quality threshold above 26.7. All segmentation parameters, including intensity thresholds and size filters, were kept consistent across all cell types (naive, EXMCs and TSCs) to ensure comparability. Segmentation results and associated statistics were exported as .csv files for downstream analysis.

Three-dimensional renderings and video animations of the confocal stacks were generated in Imaris to visualize the spatial distribution of *XIST* foci under different experimental conditions.

##### Super-resolution microscopy data analysis

To quantify the spatial distribution and clustering of *XIST* foci, a custom Python script was used to compute nearest neighbor distances (NNDs). XYZ coordinates of detected *XIST* foci were extracted from the exported .csv files (Vesicles_Type_A_Position.csv). Foci were grouped by image and cell ID, and a k-dimensional tree (KD-tree) was constructed for each group to efficiently compute Euclidean distances.

For each focus, the average distance to its five nearest neighbors (excluding itself) (NND_5) was calculated. Additionally, the number of *XIST* foci per nucleus was extracted from Cell_Number_Of_Vesicles_VesicleType=Vesicles_Type_A.csv using the same grouping strategy.

All datasets were merged into a single .xls file using image name and cell ID as keys. The merged data were further processed and visualized using custom R scripts to perform statistical analyses and generate plots.

#### RNA extraction and RT-qPCR

RNA extraction was performed referring to instructions and reagents from RNeasy® Mini Kit (QIAGEN, 74106). Briefly, cell pellets were disrupted in RLT buffer supplemented with β-mercaptoethanol and stored at -80°C until use. After adding 1 volume of 70% ethanol, total RNAs were purified on RNeasy spin columns according to the Mini Kit extraction protocol and finally eluted in RNAse-free water. Quantification of the extracted RNAs was done using the Nanodrop 2000. The DNase step was performed on 200 ng of RNA for 30 minutes at 37°C using TURBO™ DNase (Thermo Fisher Scientific). Then RNAs were reverse transcribed using the Superscript IV kit (Thermo Fisher Scientific) following the manufacturer’s recommendation. cDNAs were diluted 5 times in water, and gene expression level was assessed by real-time quantitative PCR using the Power SYBR Green Master Mix (Thermo Fisher Scientific) and ViiA-7 Real-Time PCR system (Applied Biosystems). Primers can be found in the resource table. Transcript RNA levels were normalized against *GAPDH* reference gene following the 2-ΔCt method.

#### Flow cytometry

hESCs were dissociated using Accutase (Sigma-Aldrich, A6964-100ML) into single cells by incubating for 5 minutes at 37°C. Before proceeding with the antibody staining, cells were washed twice with FACS buffer containing 1% BSA in PBS (Gibco, 10010-015). Fluorophore-conjugated antibodies were diluted at a ratio of 1:50, which is 1 μL of antibody in 50 μL of FACS buffer for around 50000 to 100000 cells, and incubated at 4°C in the dark for at least for 30 min. Cells were washed again with FACS buffer, passed through a 40 μM cell strainer (Corning, 352352340), analyzed/sorted using a BD influx or Beckman cytoflex SRT. Single-stained controls were used for compensation and setting up the precise and stringent gate in the flow cytometer. Analysis was done using the Flowjo software.

#### Bulk RNA-seq

##### Cell preparation

Cells were harvested, pelleted and stored either in Trizol Reagent (Thermo, 15596026) or TRIreagents (Sigma, T9424). RNA extraction was done using Phenol-chloroform, isopropanol, and precipitated using 70% ethanol. RNA was treated with Dnase I (Roche) to remove DNA contamination. RNA clean up was done either with 70% ethanol precipitation or with the RNeasy columns protocol (Qiagen, 74104).

##### Library Preparation

Libraries were generated using the Illumina Stranded Total RNA Prep Ligation with Ribo-Zero Plus kit (Illumina #20040525) according to the manufacturer’s recommendation and sequenced either on a NovaSeq 6000 instrument (ICGex - NGS platform, Paris, France) or on a NextSeq 2000 instrument to generate 2X100 paired-end reads.

##### Bulk RNA-seq analysis

Raw sequence reads were quality-checked using the FastQC software and cutAdapt^83^ was used to trim adapters. Reads were aligned to the human GRCh38 reference genome using STAR aligner ^84^ and deduplicated using picard. Heterozygous SNPs were found by similarly aligning H9 whole genome sequencing data from GSM1227088 ^85^ and processing it with bcftools mpileup ^86^. Expression in our bulk RNA-seq at each SNP was then determined using SNPsplit ^87^ and quantified using bedtools genomecov ^88^. Further analysis was performed in R. X-to-autosome ratios were calculated using the mean gene expression from each allele. Only SNPs with 30 or more read depth were included, as has been used previously ^89^. Allelic ratios were calculated by determining which allele was more expressed, and dividing the least expressed allele by the most expressed allele. We limited our analyses to genes known to be subject to XCI in adults ^67^, as genes known to escape from XCI were biallelic in all cell types. When multiple SNPs were available for a gene, the median allelic ratio was used.

#### CUT&RUN

CUT&RUN was performed as previously described ^90^. Briefly, 0.3 million cells per replicate were bound to 20 uL concanavalin A-coated beads (Bangs Laboratories) in binding buffer (20 mM HEPES, 10 mM KCl, 1 mM CaCl2 and 1 mM MnCl2). The beads were washed and resuspended in Dig-wash buffer (20 mM HEPES, 150 mM NaCl, 0.5 mM spermidine, 0.05% digitonin). The primary antibodies (1:50) were added to the bead slurry and rotated at RT for 1 hour. The antibodies list can be found in the resource table. The beads were washed by Dig-wash buffer and pA-Mnase fusion protein (1:400, produced by the Institut Curie Recombinant Protein Platform, 0.785mg/mL) was added and rotated at RT for 15 min. After two washes, the beads were resuspended in 150 µL Dig-wash buffer, and the MNase was activated with 2 mM CaCl2 and incubated for 30 minutes at 0 °C. MNase activity was terminated with 150 µL 2XSTOP (200 mM NaCl, 20 mM EDTA, 4 mM EGTA, 50 µg/ml RNase A and 40 µg/ml glycogen). Cleaved DNA fragments were released by incubating for 20 minutes at 37 °C, followed by centrifugation for 5 minutes at 16000g at 4 °C and collection of the supernatant from the beads on a magnetic rack. The DNA was purified by phenol:chloroform and libraries were prepared using the TruSeq ChIP Library Preparation Kit from Illumina following the manufacturer’s protocol, and sequenced on a NovaSeq 6000 instrument (ICGex - NGS platform, Paris, France) to generate 2X100 paired-end reads.

#### CUT&TAG

##### CUT&TAG Library Preparation

CT27 hTSCs were grown on iMatrix-511 (Reprocell #NP892-011) in the medium described in ^42^. Extraction of hTSC nuclei was performed as follows: cells were dissociated with Accumax (ThermoFisher Scientific #00-4666-56), washed once with PBS and resuspended in 500 µl nuclear extraction buffer consists of 20 mM HEPES-KOH pH 7.9, 10 mM KCl, 0.5 mM spermidine, 0.1% Triton X-100, 20% glycerol and 1X protease inhibitor cocktail, PIC (Roche #11697498001). Samples were centrifuged at 600g for 3 min at 4 °C in a swing rotor centrifuge, and the supernatant was discarded. The extracted nuclei were fixed with 1 ml 0.2% formaldehyde and 1X PIC in PBS for 5 min at room temperature. The fixation was stopped by adding 30 µl 2.5 M glycine, and samples were incubated for 5 min on ice. Nuclei were then centrifuged at 600 g for 3 min at 4 °C in a swing rotor centrifuge to remove the fixation solution. Finally, nuclei were resuspended in 100 µl Wash buffer (20 mM HEPES-NaOH pH 7.5, 150 mM NaCl, 0.5 mM spermidine and 1X PIC) supplemented with 0.1% BSA and 10% DMSO, and frozen slowly at -80 °C. We performed CUT&Tag as described (ref. PMID:31036827) with the following antibodies (anti-H3K27me3, CST, cat# 9733). Nuclei prepared from dissociated CT27 TSCs were immobilized to BioMag Plus Concanavalin-A-conjugated magnetic beads (ConA beads, Polysciences). Primary antibody was incubated with immobilized nuclei at 4°C overnight. Followed, secondary antibody (Guinea Pig anti-Rabbit IgG, Sigma, cat# SAB3700889) was applied and incubated at room temperature for 30 mins. Home-made pA-Tn5 was assembled with adapters and used for tagmentation of the regions bound by the target of interest. NEBnext PCR master mix (NEB, M0541) was used for tagmented fragment amplification and library preparation. We performed CUT&RUN with Naïve hPSC-converted TS-like cells using the CUTANA™ ChIC/CUT&RUN Kit (Epicypher, 14-1048). DNA libraries were prepared using the NEBNext Ultra DNA library Prep Kit (NEB, E7645S).

##### Chromatin profiling analysis

Additionally, to our CUT&RUN data, we analyzed CUT&RUN, CUT&Tag from our study and ChIP-seq ^53^. Reads were trimmed using trim_galore (0.6.5) (https://github.com/FelixKrueger/TrimGalore/) with a minimum length of 50 bp and were then mapped to the human genome (hg38) using bowtie2 (2.4.4) ^91^ with the following parameters: --local --very-sensitive --no-unal --no-mixed --no-discordant --phred33 -L 10 -X 700. Duplicated reads were removed using MarkDuplicates from picard (2.23.5) (http://broadinstitute.github.io/picard/), and aligned reads were filtered for a minimum mapping quality of 10 using samtools (1.13) ^92^. In all naive hESC and capacitated datasets, reads originating from mouse embryonic fibroblasts (MEFs) were discarded using XenofilteR package from R (4.1.1) (Kluin et al., 2018). To create the correlation heatmap, mean read depth per 10kb windows of the X chromosome were obtained using mosdepth (0.2.3) ^93^ and then processed on R (4.3.2) to be normalized by library size and used to compute Spearman’s rank correlation coefficient and for the 500 most variable regions for each histone mark. To create all the other figures, aligned reads corresponding to replicates were merged and downsampled to the smallest library’s read count (26,5M) using samtools (1.13) ^92^. BigWig track files displaying the log2 of histone mark enrichment ratio over IgG were generated using bigwigCompare from deeptools (3.5.0) ^94^ with the --operation log2 parameter, on histone mark and IgG BigWig files obtained with bamCoverage from deeptools (3.5.0) ^94^ using the following parameters: --normalizeUsing BPM --binSize 20 --smoothLength 40. We used samtools idxstats (1.13) ^92^ to obtain the total number of reads mapping to each chromosome. We calculated occupancy percentages by determining binding sites by peak calling using macs2 (2.2.7.1) ^95^ with the following parameters: -f BAMPE -g 2913022398 --keep-dup ‘all’ --broad --broad-cutoff 0.1 -- nolambda, then merging peaks less than 10kb apart using bedtools (2.30.0) ^88^ and finally computing in R (4.3.2) the cumulative width of all peaks over the chromosome size. To analyze the annotations covered by the histone marks, we used bulk RNA-seq data of naive hESCs ^26^, EXMCs and TSCs (this study) and the cDNA and ncRNA (GRCh38) sequences downloaded from ENSEMBL (www.ensembl.org) select the most expressed isoform for each X-linked gene in each cell type using kallisto (0.46.2) ^96^, and constructed an annotation file for each cell type from these results. Gene bodies were defined as the sequences between the Transcription Start Site (TSS) and Transcription End Site (TES) given in kallisto outputs, promoters as the 1kb region preceding the TSS, the ‘upstream’ regions as the 4kb regions preceding promoters, predicted enhancers were downloaded from the FANTOM5 consortium collection hg38_v9 ^97^, and the remaining was defined as intergenic. An annotation was considered covered by a histone mark if 75% of the annotation sequence overlapped with a peak.

#### MeD-seq

MeD-seq assays were essentially performed as previously described ^64^. Briefly, DNA from naive hESCs, EXMCs and TSCs were extracted using the column DNeasy kit (Qiagen, 69504) following the manufacturer’s recommendations. Then, 10 μl genomic DNA (input 90 ng) from naive hESCs, EXMCs and TSCs were digested with LpnPI (New England Biolabs) generating 32 bp fragments around the fully methylated recognition site containing a CpG. These short DNA fragments were further processed using the ThruPlex DNA–seq 96D kit (Rubicon Genomics Ann Arbor). Stem-loop adapters were blunt-end ligated to repair input DNA and amplified to include dual indexed barcodes using a high-fidelity polymerase to generate an indexed Illumina NGS library. The amplified end product was purified on a Pippin HT system with 3% agarose gel cassettes (Sage Science). Multiplexed samples were sequenced on Illumina NextSeq2000 systems for paired-end read of 50 bp according to the manufacturer’s instructions. Dual indexed samples were de-multiplexed using bcl2fastq software (Illumina).

##### MeD-seq analysis

Reads were trimmed using trim_galore (0.6.5) (https://github.com/FelixKrueger/TrimGalore/) and were then mapped to the human genome (hg38) and the mouse genome (mm10) using bowtie2 (2.4.4) ^91^ with the following parameters: --local --very-sensitive --no-unal --phred33 -L 10. BAM files were filtered to discard reads with mapping quality lower than 10 using samtools (1.13) ^92^ and reads originating from mouse embryonic fibroblasts using XenofilteR package from R (4.1.1) ^98^. BigWig files were generated using bamCoverage from deeptools (3.5.0) ^94^ with the following parameters: normalizeUsing=“None”, binSize=20, smoothLength=40. CpG-island annotations were downloaded from ENSEMBL (www.ensembl.org). Coverage scores were then computed in R (4.3.2) for each CpG located on the X chromosome, as log2 (x + 1), x being the number of reads per bin. Coverage scores were finally summed across the entire X chromosome or within specific genomic features. Gene bodies were defined as the sequences between the Transcription Start Site (TSS) and Transcription End Site (TES) based on Ensembl annotation release 90 for GRCh38.p10 (www.ensembl.org), TSS regions as ± 1 kb around the TSS, and predicted enhancers were downloaded from the FANTOM5 consortium collection (hg38_v9) ^97^.

#### scRNA-seq

Cells were dissociated with Accutase (Stemcell Technologies, 07920) for 6 minutes (for naive hESCs day 0) or with TrypLE for 15 minutes (for day 4 and day 8 of differentiation) at 37°C. Dissociation was stopped by adding DMEM and cells were pelleted and resuspended in PXGL or ASECRiAV. Feeder depletion was performed by placing cell suspensions on a gelatin pre-coated plate for 35-45 minutes. Cell counts were calculated using LunaFL (Logos Biosystems) and Propidium iodide stain (Logos, F23001). At least 100000 cells were used for further steps. Samples were fixed and permeabilized using Chromium NEXT GEM Single Cell Fixed RNA Sample (10x Genomics, #1000414). Briefly, at each time point, cell pellets were resuspended in PFA and permeabilization reagents for 10 hours at 4°C. Suspensions were centrifuged and resuspended in quenching buffer, enhancer reagent, and glycerol and kept at -80°C before further manipulation. Probes were hybridized, to add barcodes specific to each sample. Samples from each day were pooled together to obtain 20000 cells per sample, unbound probes were then washed off. Gel beads in emulsion (GEM) were then generated to contain a unique barcoded gel bead, one cell and reaction enzyme and reagents in each droplet (Chromium, PN 1000475). Ligation, extension, clean up and library construction were then conducted according to the manufacturer’s recommendation. Pooled libraries were then sequenced on an Illumina Nextseq 2000 sequencing platform using a P3 100 cycles kit to a minimum sequencing depth of 10000 reads per cell using reads lengths of 28 bp read 1, 10 bp double indexes and 90 bp read 2.

##### Single-cell RNA-seq analysis

Single cell RNA-seq analysis was done at multiple time points in WT or XIST iKD cells as shown in Figure 4A and as explained in ^44^ with modifications. Raw sequence reads were quality-checked using the FastQC software. The CellRanger version 7.0.1 was used to process, align and summarize unique molecular identifier (UMI) counts against the 10X Genomics pre-built human GRCh38 reference genome dataset (2020-A, July 7, 2020). Downstream analyses were performed in R using Seurat (v4.0.1) ^99^. Cell with good sequencing reads were retained by adjusting the number of counts per cell (nCount_RNA) and the number of mapped genes per cell (nFeature_RNA) to only keep cells that were mostly mapped to the human GRCh38 (hg38) genome (nCount _RNA > 2200, nFeature_RNA >1500). Cells with more than 25% of mitochondrial counts were filtered out. After quality control, we obtained data for 64479 cells with 28091 cells on day 0, 26650 cells on day 4 and 9738 cells on day 8. The count matrix was normalized with Seurat global-scaling normalization method “LogNormalize’’ that normalizes the feature expression measurements for each cell by the total expression, multiplies this by a 10000-scale factor, and log-transforms the result. Cells were annotated based on gene expression. Differential expression testing was performed with the FindMarkers function in Seurat based on the non-parametric Wilcoxon rank sum test applying the logFC threshold of averaged log_2_ FC > 0.25. A graph-based cell clustering approach was used to cluster cells with the FindClusters function in Seurat. Allelic gene expression was calculated using the scLinaX package for R ^65^. Heterozygous SNPs were first identified using bcftools, and SNPs were filtered to only include those with >20 total reads, and >5 reads from each SNP across all cell types and time points. Vartrix was used to obtain SNP depth per allele for each cell and ANNOVAR^100^ was used to annotate and filter SNPs with less than 0.01 minor allele frequency in the gnomAD v4.1 database ^101^. The allelic data was then processed further in R using the scLinaX pipeline. We used the default XCI status annotations from ^66^.

#### Quantification and statistical analysis

Statistical tests and data processing were performed in R (v4.0.3). All statistical methods and test values used can be found in the relevant result section, figures, and figure legends. In bulk RNA-seq, CUT&RUN, and Med-seq experiments, two replicates were used. In scRNA-seq experiments, one replicate was included per time point, and the number of cells per condition is indicated in **Table S1**. During the processing step of the scRNA-seq cells were excluded with nCounts RNA<2200, nFeature RNA<1500, and >25% of mitochondrial reads. Differential gene expression analysis was performed using the Seurat function FindMarkers using cutoffs of adjusted p-value<0.05 and log2 fold change>0.5.

All the experiments were repeated at least two or three times, and with cells with different genetic backgrounds, with exceptions. Sequencing experiments were done once, with or without replicates. Aside from the scRNA-seq experiment, all other *XIST* iKD experiments, including IF, FISH, differentiation, flow cytometry, and brightfield imaging, were performed using three independent XIST iKD clones, with the exception of Annexin V flow cytometry data, where 2 clones were used. For the *XIST* iKD, three independent clones were used for all experiments related to Figures **4B-4E and 5F-5G**. Experiments were repeated three times for **1C-1H and 3A-3F,** once for **1B, 2A-2H, 4F-4L, and 5F-5G,** and two times for all other figures. Experiments were not blinded. No data were excluded with the exception of cell filtering described in the scRNA-seq analysis as well as in this section above. Specifically, a logistic regression test was used for **Figures 1D, 1F, 1H, 3B, 3D, 3F, 3K, 3M, and S5I;** a chi-squared test for **Figures 1J, 4E, S6C, and S6G**; a t-test for **Figures S1R, 3I, S4C, 4D, 5A, 5B, S5E-S5G, and S6C-S6D, S6G;** and a Wilcoxon rank-sum test for **Figures S1I-S1J, S4J-S4K, and S4D-S4E.**

## ACKNOWLEDGEMENTS

We thank Dr. Rudolf Jaenisch and Dr. Thorold W. Theunissen for the WIBR2-29M-GP26-TN9 hESC line. We are grateful to the Vlaams Supercomputer Center Leuven (https://www.vscentrum.be) for computing, in particular Jan Ooghe; the VIB/KU Leuven Center for Brain and Disease Research, in particular to Kristofer Davie. We are also grateful to the Genomics Core Leuven (http://genomicscore.be) for high-throughput sequencing and 10X Chromium Controller, and in particular Greet Bervoets for helpful feedback. We are also grateful to the whole KU Leuven FACS core team for providing the facility and especially Reena Chinaraj who helped us during flow cytometry experiments. We thank Pablo Hernandez-Varas, Charlotte Cresens, and the whole VIB bioimaging core, KU Leuven team for providing training and valuable resources for confocal and 3D-SIM imaging. We are also grateful to the Statistics and Data Science Department, KU Leuven for providing valuable suggestions for the statistical data analysis used in the paper. The Pasque lab is part of the Leuven Single-Cell Omics Institute (LISCO) and Leuven Stem Cell Institute (SCIL). Research in the Pasque laboratory is supported by The Research Foundation–Flanders (FWO; Odysseus Return Grant G0F7716N to V.P.; FWO grants G0C9320N and G0B4420N to V.P.), the KU Leuven Research Fund (V.P. C1 grants C14/21/19 to V.P. and project financing), FWO PhD fellowships to S.K.T. (1S75720N), T.X.A.P.(11N3122N), R.N.A. (11L0722N) and junior postdoctoral FWO fellowship to B.P.B (1263323N).

We thank Mauro Calabrese for giving the vectors necessary for the piggy-BAC system (PB_rtTA_BsmBI vector and PB_tre_dCas9_KRAB). We thank Epigenomic, Microscopy, Vectorology and BIBS Platforms, all hosted in UMR7216 Epigenetic and Cell Fate, for technical advice and access to instruments. We thank the ICGex NGS platform of the Institut Curie supported by the grants ANR-10-EQPX-03 (Equipex) and ANR-10-INBS-09-08 (France Génomique Consortium) from the Agence Nationale de la Recherche (“Investissements d’Avenir” program), by the ITMO-Cancer Aviesan (Plan Cancer III) and by the SiRIC-Curie program (SiRIC Grant INCa-DGOS-465 and INCa-DGOSInserm_12554) for the high-throughput sequencing. We thank the Bioinformatics platform of the Institut Curie for data management, quality control and primary analysis. We acknowledge Brianna Rodgers for the preparation of the 10x libraries, thanks to the funding from the Labex “Who am I” Single Cell Initiative. We thank Brigitte Izac from the Genomic facility of Institute Cochin for her precious help in the production of 10X libraries and their sequencing. This project was funded by Agence Nationale pour la Recherche [ANR-14-CE10-0017 to C.R.]; European Research Council (ERC) under the European Union’s Horizon 2020 research and innovation program [101020423]; LabEx ‘Who Am I?’ [ANR-11-LABX-0071]; Université de Paris IdEx [ANR-18-IDEX-0001] funded by the French Government through its ‘Investments for the Future’ program. L.C. was supported by fellowships from the French Ministry of Education and Research and LabEx ‘Who Am I? (ANR-11-LABX-0071) and Idex (ANR-18-IDEX-0001). E.C. was supported by fellowships from the French Ministry of Education and Research and from the French Medical Research Foundation (FRM).

Research in the Rugg-Gunn lab is supported by grants from the BBSRC (BBS/E/B/000C0521; Core Capability Grant), MRC (MR/T011769/1; MR/V02969X/1) and Wellcome (215116/Z/18/Z; 225839/Z/22/Z).

Research in the Niakan lab is supported by Wellcome grants (221856/Z/20/Z; 215116/Z/18/Z).

## AUTHOR CONTRIBUTIONS

Conceptualization: A.P., L.C., J.F.O., C.R., and V.P.; data curation: A.P., L.C.; Formal analysis: A.P., L.C., J.F.O., C.R., and V.P.; Funding acquisition: C.R., and V.P.; Investigation (omics experiments): A.P., L.C., B.P.B., J.B., C.A., G.C., R.B., S.K., S.K.T., T.X.A.P., R.N.A., J.G., Y.W., D.S., K.K.N., P.J.R.G.; Investigation (stem cell experiments): A.P., L.C., R.S.T., J.FO., C.L., J.B., S.N., E.C., M.M.; Methodology: A.P., L.C., J.F.O., C.R., and V.P.; Project administration: A.P., L.C., J.F.O., C.R., and V.P.; Resources: C.R., and V.P.; Supervision: J.FO., C.R., and V.P.; Validation: A.P., L.C., J.F.O., C.R., and V.P. Visualization: A.P., L.C., J.F.O., C.R., and V.P.; writing, reviewing, and editing of the manuscript: A.P., L.C., B.P.B., J.B., J.F.O., C.R., and V.P.

